# Mapping neurotransmitter systems to the structural and functional organization of the human neocortex

**DOI:** 10.1101/2021.10.28.466336

**Authors:** Justine Y. Hansen, Golia Shafiei, Ross D. Markello, Kelly Smart, Sylvia M. L. Cox, Martin Nørgaard, Vincent Beliveau, Yanjun Wu, Jean-Dominique Gallezot, Étienne Aumont, Stijn Servaes, Stephanie G. Scala, Jonathan M. DuBois, Gabriel Wainstein, Gleb Bezgin, Thomas Funck, Taylor W. Schmitz, R. Nathan Spreng, Marian Galovic, Matthias J. Koepp, John S. Duncan, Jonathan P. Coles, Tim D. Fryer, Franklin I. Aigbirhio, Colm J. McGinnity, Alexander Hammers, Jean-Paul Soucy, Sylvain Baillet, Synthia Guimond, Jarmo Hietala, Marc-André Bédard, Marco Leyton, Eliane Kobayashi, Pedro Rosa-Neto, Melanie Ganz, Gitte M. Knudsen, Nicola Palomero-Gallagher, James M. Shine, Richard E. Carson, Lauri Tuominen, Alain Dagher, Bratislav Misic

## Abstract

Neurotransmitter receptors support the propagation of signals in the human brain. How receptor systems are situated within macroscale neuroanatomy and how they shape emergent function remains poorly understood, and there exists no comprehensive atlas of receptors. Here we collate positron emission tomography data from >1 200 healthy individuals to construct a whole-brain 3-D normative atlas of 19 receptors and transporters across 9 different neurotransmitter systems. We find that receptor profiles align with structural connectivity and mediate function, including neurophysiological oscillatory dynamics and resting state hemodynamic functional connectivity. Using the Neurosynth cognitive atlas, we uncover a topographic gradient of overlapping receptor distributions that separates extrinsic and intrinsic psychological processes. Finally, we find both expected and novel associations between receptor distributions and cortical thinning patterns across 13 disorders. We replicate all findings in an independently collected autoradiography dataset. This work demonstrates how chemoarchitecture shapes brain structure and function, providing a new direction for studying multi-scale brain organization.

## Introduction

The brain is a complex system that integrates signals across spatial and temporal scales to support cognition and behaviour. The key neural signalling molecule is the neurotransmitter: chemical agents that relay messages across synapses. While neurotransmitters carry the message, neurotransmitter receptors act as ears that cover the cellular membrane, determining how the postsynaptic neuron will respond. By modulating the excitability and firing rate of the cell, neurotransmitter receptors effectively mediate the transfer and propagation of electrical impulses. As such, neurotransmitter receptors drive synaptic plasticity, modify neural states, and ultimately shape network-wide communication [121, 123, 137].

How spatial distributions of neurotransmitter receptors relate to brain structure and shape brain function at the system level remains unknown. Recent technological advances allow for high resolution reconstructions of the brain’s wiring patterns. These wiring patterns display non-trivial architectural features including specialized network modules that support the segregation of information [128], as well as densely interconnected hub regions that are thought to support the integration of information [127, 138]. The spatial arrangement of neurotransmitter receptors on this network presumably guides the flow of information and the emergence of cognitive function. Therefore, understanding the link between structure and function is inherently incomplete without a comprehensive map of the chemoarchitecture of the brain [63, 90, 154, 156].

A primary obstacle to studying the relative density distributions of receptors across multiple neurotransmitter systems is the lack of comprehensive openly accessible datasets. An important exception is the autoradiography dataset of 15 neurotransmitter receptors and receptor binding sites, collected in three post-mortem brains [154]. However, these autoradiographs are only available in 44 cytoarchitectonically defined cortical areas. Alternatively, positron emission tomography (PET) can estimate *in vivo* receptor concentrations across the whole brain. Despite the relative ease of mapping receptor densities using PET, there are nonetheless difficulties in constructing a comprehensive PET dataset of neurotransmitter receptors. Due to the radioactivity of the injected PET tracer, mapping multiple different receptors in the same individual is not considered a safe practice. Combined with the fact that PET image acquisition is relatively expensive, cohorts of control subjects are small and typically only include one or two tracers. Therefore, constructing a comprehensive atlas of neurotransmitter receptor densities across the brain requires extensive data sharing efforts from multiple research groups [12, 30, 62, 71, 84, 86].

Here we curate and share an atlas of PET-derived whole-brain neurotransmitter receptor maps from 19 unique neurotransmitter receptors, receptor binding sites, and transporters, across 9 different neurotransmitter systems and over 1 200 healthy individuals, available at https://github.com/netneurolab/hansen_receptors. We use multiple imaging modalities to comprehensively situate neurotransmitter receptor densities within microscale and macroscale neural architectures. Using diffusion weighted MRI and functional MRI, we show that neurotransmitter receptor densities follow the organizational principles of the brain’s structural and functional connectomes. Moreover, we find that neurotransmitter receptor densities shape magnetoencephalography (MEG)-derived oscillatory neural dynamics. To determine how neurotransmitter receptor distributions affect cognition and disease, we map receptor densities to meta-analytic (Neurosynth-derived) functional activations, where we uncover a spatially covarying axis of neuromodulators and mood-related processes. Next, we link receptor distributions to ENIGMA-derived patterns of cortical atrophy across 13 neurological, psychiatric, and neurodevelopmental disorders, uncovering specific receptor-disorder links. We validate our findings and extend the scope of the investigation to additional receptors using an independently collected autoradiography neurotransmitter receptor dataset [155]. Altogether we demonstrate that, across spatial and temporal scales, chemoarchitecture consistently plays a key role in brain function.

## Results

A comprehensive cortical profile of neurotransmitter receptor densities was constructed by collating PET images from a total of 19 different neurotransmitter receptors, transporters, and receptor binding sites, across 9 different neurotransmitter systems, including dopamine, norepinephrine, serotonin, acetylcholine, glutamate, GABA, histamine, cannabinoid, and opioid (Fig. 1). All PET images are acquired in healthy participants (see Table 1 for a complete list of receptors and transporters, corresponding PET tracers, ages, and number of participants). Each PET tracer map was processed according to the best practice for the radioligand; for detailed acquisition and processing protocols see the publications listed in Table 1. A group-average tracer map was constructed across participants within each study. To mitigate variation in image acquisition and preprocessing, and to ease biological interpretability, all PET tracer maps were parcellated into the same 68 cortical regions and z-scored [26]. After parcellating and normalizing the data, maps from different studies of the same tracer were averaged together (Fig. S1 shows consistencies across studies). In total, we present tracer maps for 19 unique neurotransmitter receptors and transporters from a combined total of 1239 healthy participants, resulting in a 68 × 19 matrix of relative neurotransmitter receptor/transporter densities. Finally, we repeat all analyses in an independently collected autoradiography dataset of 15 neurotransmitter receptors (Table S1; [155]), and across alternative brain parcellations [18].

**Figure 1.**
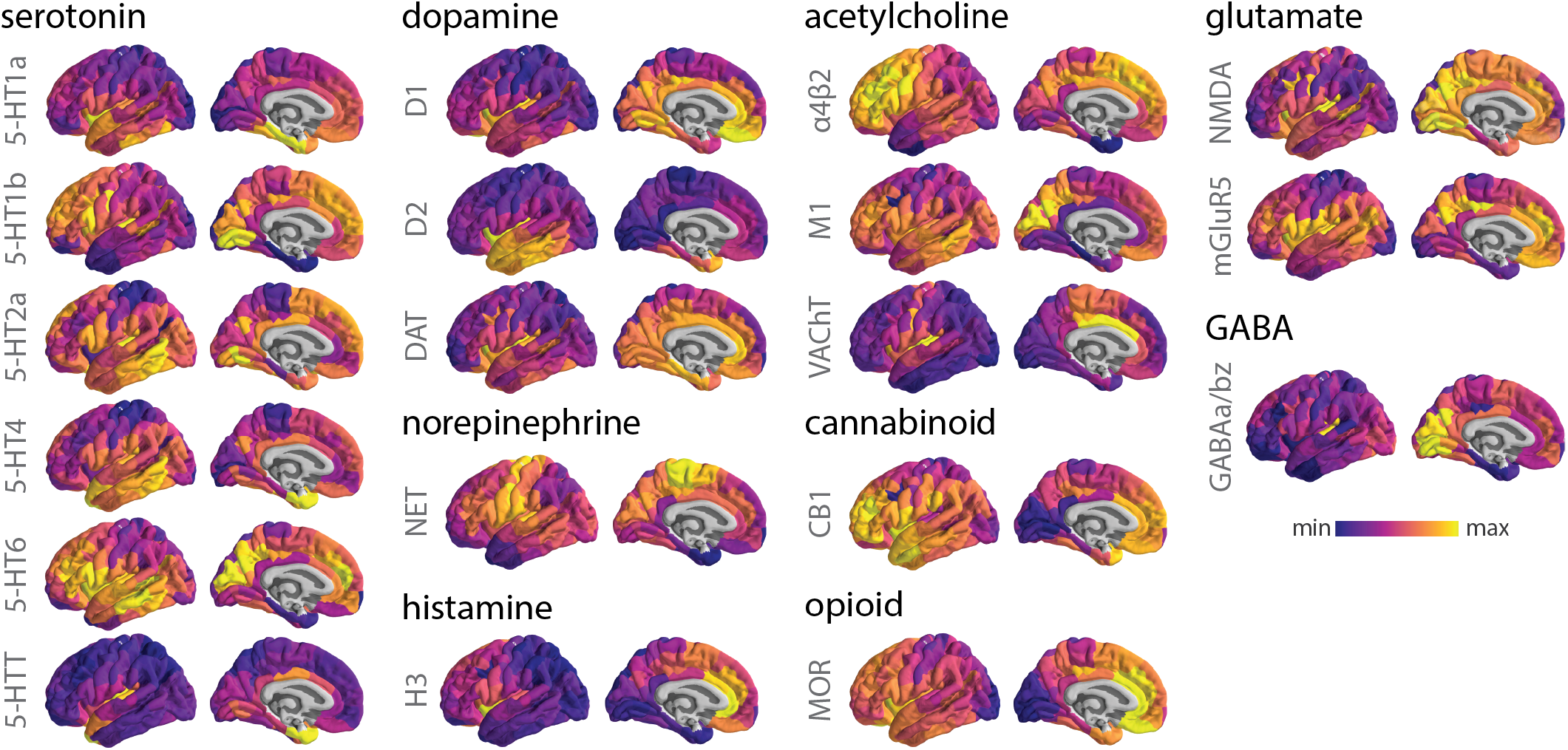
PET images of neurotransmitter receptors and transporters. PET tracer images were collated and averaged to produce mean receptor distribution maps of 19 different neurotransmitter receptors and transporters across 9 different neurotransmitter systems and a combined total of over 1 200 healthy participants.

**TABLE 1.** Neurotransmitter receptors and transporters included in analyses. BP_ND_ = non-displaceable binding potential; V_T_ = tracer distribution volume; B_max_ = density (pmol/ml) converted from binding potential (5-HT) or distributional volume (GABA) using autoradiography-derived densities; SUVR = standard uptake value ratio. Neurotransmitter receptor maps without citations refer to unpublished data. In those cases, contact information for the study PI is provided in Table S3. Table S3 also includes more extensive methodological details such as PET camera, number of males and females, modelling method, reference region, scan length, and modelling notes. Asterisks indicate transporters.

| Receptor/<br>transporter | Neurotransmitter | Tracer | Measure | *N* | Age | References |
| --- | --- | --- | --- | --- | --- | --- |
| D<sub>1</sub> | dopamine | [<sup>11</sup>C]SCH23390 | BP<sub>ND</sub> | 13 | 33 ± 13 | Kaller et al., 2017 [58] |
| D<sub>2</sub> | dopamine | [<sup>11</sup>C]FLB-457 | BP<sub>ND</sub> | 37 | 48.4 ± 16.9 | Smith et al., 2019 [109, 126] |
| D<sub>2</sub> | dopamine | [<sup>11</sup>C]FLB-457 | BP<sub>ND</sub> | 55 | 32.5 ± 9.7 | Sandiego et al., 2015 [109, 110, 124, 126, 153] |
| DAT* | dopamine | [<sup>123</sup>I]-FP-CIT | SUVR | 174 | 61 ± 11 | Dukart et al., 2018 [29] |
| NET* | norepinephrine | [<sup>11</sup>C]MRB | BP<sub>ND</sub> | 77 | 33.4 ± 9.2 | Ding et al., 2010 [11, 19, 27, 108] |
| 5-HT<sub>1A</sub> | serotonin | [<sup>11</sup>C]WAY-100635 | BP<sub>ND</sub> | 36 | 26.3 ± 5.2 | Savli et al., 2012 [113] |
| 5-HT<sub>1B</sub> | serotonin | [<sup>11</sup>C]P943 | BP<sub>ND</sub> | 65 | 33.7 ± 9.7 | Gallezot et al., 2010 [9, 39, 72, 80, 81, 94, 111] |
| 5-HT<sub>1B</sub> | serotonin | [<sup>11</sup>C]P943 | BP<sub>ND</sub> | 23 | 28.7 ± 7.0 | Savli et al., 2012 [113] |
| 5-HT<sub>2A</sub> | serotonin | [<sup>11</sup>C]Cimbi-36 | B<sub>max</sub> | 29 | 22.6 ± 2.7 | Beliveau et al., 2017 [12] |
| 5-HT<sub>4</sub> | serotonin | [<sup>11</sup>C]SB207145 | B<sub>max</sub> | 59 | 25.9 ± 5.3 | Beliveau et al., 2017 [12] |
| 5-HT<sub>6</sub> | serotonin | [<sup>11</sup>C]GSK215083 | BP<sub>ND</sub> | 30 | 36.6 ± 9.0 | Radhakrishnan et al., 2018 [98, 99] |
| 5-HTT* | serotonin | [<sup>11</sup>C]DASB | B<sub>max</sub> | 100 | 25.1 ± 5.8 | Beliveau et al., 2017 [12] |
| α<sub>4</sub>β<sub>2</sub> | acetylcholine | [<sup>18</sup>F]flubatine | V<sub>T</sub> | 30 | 33.5 ± 10.7 | Hillmer et al., 2016 [8, 52] |
| M<sub>1</sub> | acetylcholine | [<sup>11</sup>C]LSN3172176 | BP<sub>ND</sub> | 24 | 40.5 ± 11.7 | Naganawa et al., 2021 [82] |
| VACHT* | acetylcholine | [<sup>18</sup>F]FEOBV | SUVR | 4 | 37 ± 10.2 | PI: Lauri Tuominen & Synthia Guimond |
| VACHT* | acetylcholine | [<sup>18</sup>F]FEOBV | SUVR | 18 | 66.8 ± 6.8 | Aghourian et al., 2017 [1] |
| VACHT* | acetylcholine | [<sup>18</sup>F]FEOBV | SUVR | 5 | 68.3 ± 3.1 | Bedard et al., 2019 [10] |
| VACHT* | acetylcholine | [<sup>18</sup>F]FEOBV | SUVR | 3 | 66.6 ± 0.94 | PI: Taylor W. Schmitz & R. Nathan Spreng |
| NMDA | glutamate | [<sup>18</sup>F]GE-179 | V<sub>T</sub> | 29 | 40.9 ± 12.7 | Galovic et al., 2021 [41, 42, 73] |
| mGluR<sub>5</sub> | glutamate | [<sup>11</sup>C]ABP688 | BP<sub>ND</sub> | 73 | 19.9 ± 3.04 | Smart et al., 2019 [125] |
| mGluR<sub>5</sub> | glutamate | [<sup>11</sup>C]ABP688 | BP<sub>ND</sub> | 22 | 67.9 ± 9.6 | PI: Pedro Rosa-Neto & Eliane Kobayashi |
| mGluR<sub>5</sub> | glutamate | [<sup>11</sup>C]ABP688 | BP<sub>ND</sub> | 28 | 33.1 ± 11.2 | DuBois et al., 2016 [28] |
| GABA<sub>A/BZ</sub> | GABA | [<sup>11</sup>C]flumazenil | B<sub>max</sub> | 16 | 26.6 ± 8 | Nørgaard et al., 2021 [84] |
| H<sub>3</sub> | histamine | [<sup>11</sup>C]GSK189254 | V<sub>T</sub> | 8 | 31.7 ± 9.0 | Gallezot et al., 2017 [40] |
| CB<sub>1</sub> | cannabinoid | [<sup>11</sup>C]OMAR | V<sub>T</sub> | 77 | 30.0 ± 8.9 | Normandin et al., 2015 [31, 83, 87, 100] |
| MOR | opioid | [<sup>11</sup>C]carfentanil | BP<sub>ND</sub> | 204 | 32.3 ± 10.8 | Kantonen et al., 2020 [59] |

### Receptor distributions reflect structural and functional organization

Consistent with previous reports, we find that mean density across all neurotransmitter receptors and transporters are distributed heterogeneously across the cortex, with enrichment in insular and cingulate regions (Fig. 2a; [48, 155]). In addition, receptor/transporter maps are generally positively correlated with one another (Fig. 2b). To quantify the potential for two brain regions to be similarly modulated by endogenous or exogenous input, we compute the correlation of receptor/transporter fingerprints between pairs of brain regions (Fig. 2c, d). Hereafter, we refer to this quantity as “receptor similarity”, analogous to other commonly used measures of inter-regional attribute similarity including anatomical covariance [34], morphometric similarity [118], gene coexpression [5, 37, 101], temporal profile similarity [120], and microstructural similarity [92]. We confirm that no single receptor or transporter exerts undue influence on the receptor similarity matrix (see *Sensitivity and robustness analyses*).

**Figure 2.**
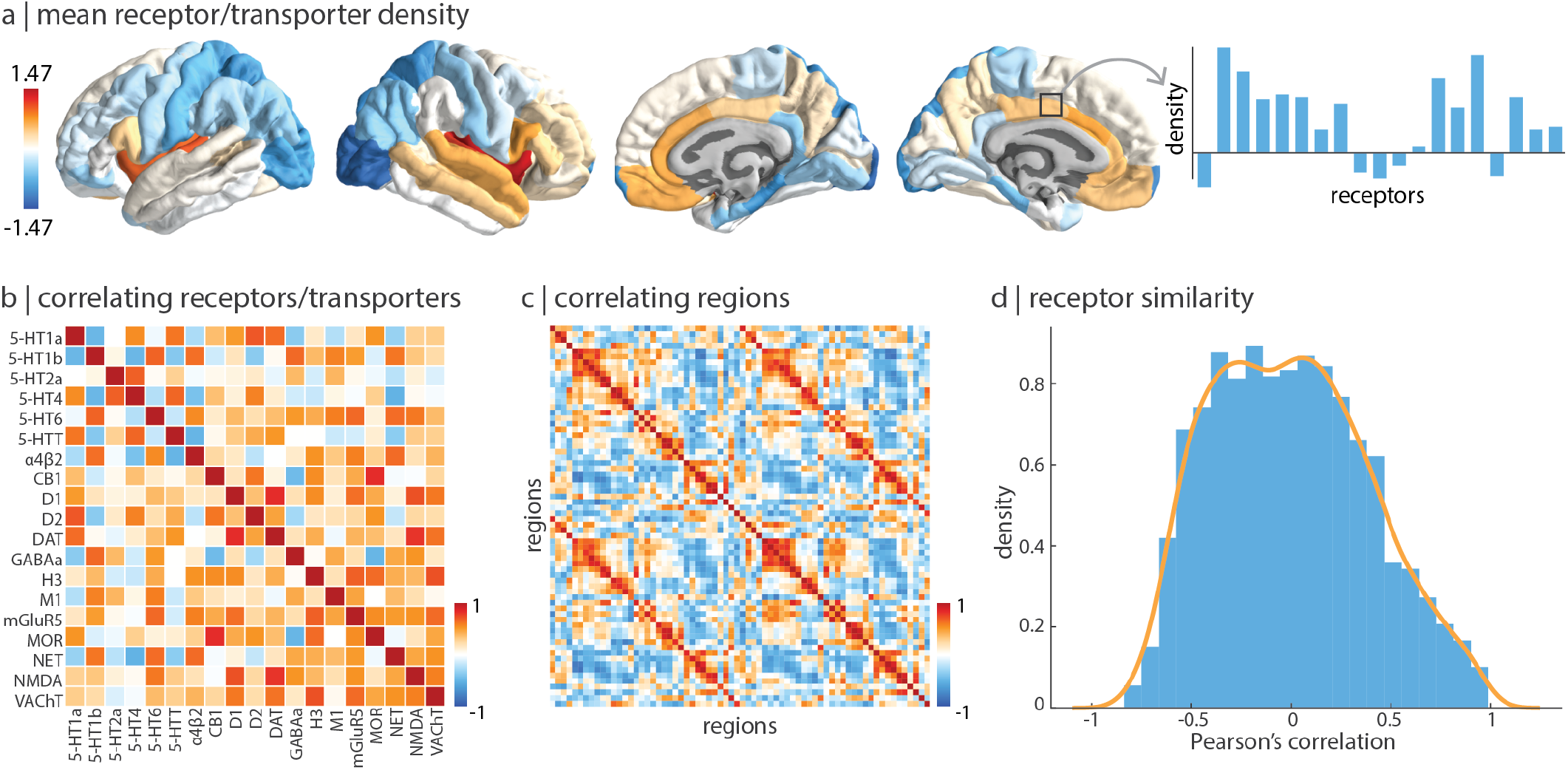
Constructing a cortical neurotransmitter receptor and transporter atlas. PET maps for 19 different neurotransmitter receptors and transporters were z-scored and collated into a single neurotransmitter receptor atlas. (a) The mean neurotransmitter receptor/transporter density across the cortex reveals that neurotransmitter receptors and transporters are enriched in insular and limbic regions. (b) Pearson’s correlation between pairs of receptor/transporter distributions across 68 brain regions. (c) The “receptor similarity” matrix is constructed by correlating (Pearson’s *r*) receptor/transporter fingerprints between pairs of brain regions. (d) The distribution of receptor similarity values.

Using group-consensus structural and resting-state functional connectomes from 70 individuals (see *Methods* for details), we show that neurotransmitter receptor organization reflects structural and functional connectivity. Specifically, we find that receptor similarity is greater between pairs of brain regions that are structurally connected, suggesting that anatomically connected areas are likely to be co-modulated (Fig. 3a). To ensure the observed relationship between structural connections and receptor similarity is not due to spatial proximity or network topography, we assessed significance against density-, degree- and edge length-preserving surrogate structural connectivity matrices (*p* = 0.0001, 10 000 repetitions [13]). Additionally, we find that receptor similarity decays exponentially with Euclidean distance a finding that has been reported for correlated gene expression [35], temporal similarity [120], structural connectivity [47, 54, 103, 107], and functional connectivity [106].

**Figure 3.**
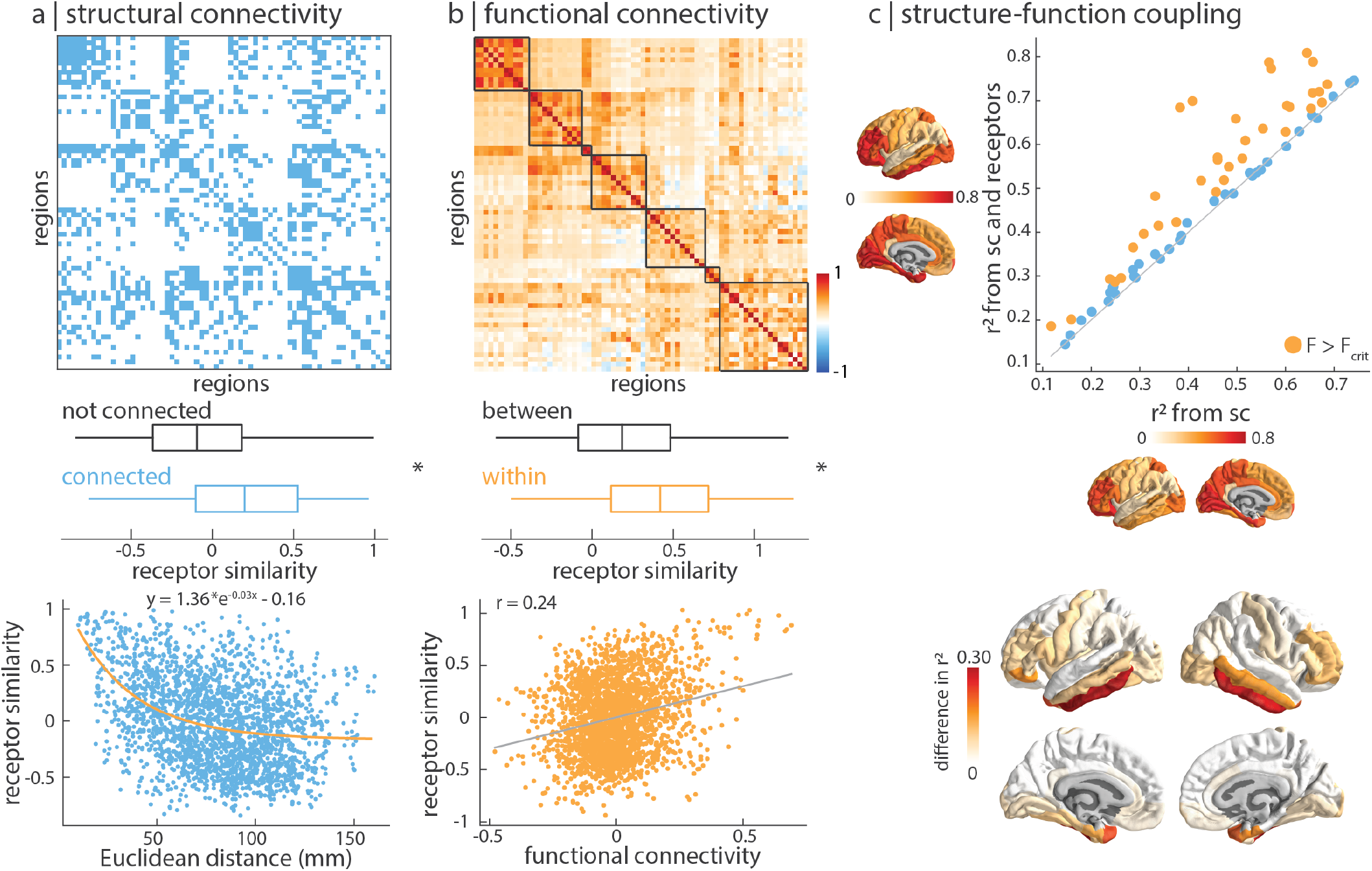
Receptor distributions reflect structural and functional organization. (a) Top: group-consensus structural connectivity matrix. Middle: receptor similarity is significantly greater between regions that are physically connected, against distance- and edge length-preserving null structural connectivity matrices (*p* = 0.0001; [13]). Bottom: receptor similarity decreases exponentially with Euclidean distance. (b) Top: group-average functional connectivity matrix, ordered by intrinsic networks [152]. Middle: receptor similarity is significantly greater within regions in the same functional network (*p*_spin_ = 0.016). Bottom: receptor similarity is positively correlated with functional connectivity (*r* = 0.24, *p* = 7 × 10^−33^). (c) Regional structure-function coupling was computed as the fit (*R*^2^) between measures of structural connectivity and functional connectivity. Top: structure-function coupling at each brain region is plotted when receptor similarity is excluded (*x*-axis) and included (*y*-axis) in the model. Yellow points indicate brain regions where receptor information significantly augments structure-function coupling (F > F_critical_). Bottom: the difference in adjusted *R*^2^ when receptor similarity is and isn’t included in the regression model. Asterisks in panels (a) and (b) denote significance. Boxplots in panels (a) and (b) represent the 1st, 2nd (median) and 3rd quartiles, whiskers represent the non-outlier end-points of the distribution, and diamonds represent outliers.

Likewise, receptor similarity is significantly greater between brain regions that are within the same intrinsic networks than between different intrinsic networks, according to the Yeo-Krienen 7-network classification (*p*_spin_ = 0.016, 10 000 repetitions, Fig. 3b [152]). This suggests that areas that are in the same cognitive system tend to have similar receptor profiles [154]. Significance was assessed non-parametrically by permuting the intrinsic network affiliations while preserving spatial autocorrelation (“spin test”; [3, 70]). We also find that receptor similarity is significantly correlated with functional connectivity, after regressing Euclidean distance from both matrices (*r* = 0.24, *p* = 7 × 10^−33^). In other words, we observe that brain regions with similar receptor and transporter composition show greater functional co-activation. Collectively, these results demonstrate that receptor profiles are systematically aligned with patterns of structural and functional connectivity above and beyond spatial proximity, consistent with the notion that receptor profiles guide inter-regional signaling.

Since neurotransmitter receptor and transporter distributions are organized according to structural and functional architectures, we next asked whether receptor/transporter distributions might augment the coupling between brain structure and function. To quantify structure-function coupling, we used a method previously developed and validated [143] in which regional structure-function coupling is defined as the adjusted *R*^2^ of a multilinear regression model that fits measures of the structural connectome to functional connectivity. We then included receptor similarity as an independent variable, to assess how receptor information changes structure-function coupling. We find that receptor information significantly augments the coupling between brain structure and function in multiple regions, especially the inferior temporal and dorsolateral prefrontal cortex (F*>*F_critical_; Fig. 3c).

### Receptor profiles shape oscillatory neural dynamics

Given that neurotransmitter receptors modulate the firing rates of neurons and therefore population activity, we sought to relate the cortical patterning of neurotransmitter receptors to neural oscillations [122]. We used MEG power spectra across six canonical frequency bands from 33 unrelated participants in the Human Connectome Project (see *Methods* for details; [45, 119, 140]). We fit a multiple linear regression model that predicts the cortical power distribution of each frequency band from neurotransmitter receptor and transporter densities. We then cross-validated the model using a distance-dependent method that was previously developed inhouse (Fig. S2; see *Methods* for details [49]). We found a close fit between receptor densities and MEG-derived power (0.85 ≤ 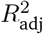 ≤ 0.94; Fig. 4a), suggesting that overlapping spatial topographies of multiple neurotransmitter systems may ultimately manifest as coherent oscillatory patterns.

**Figure 4.**
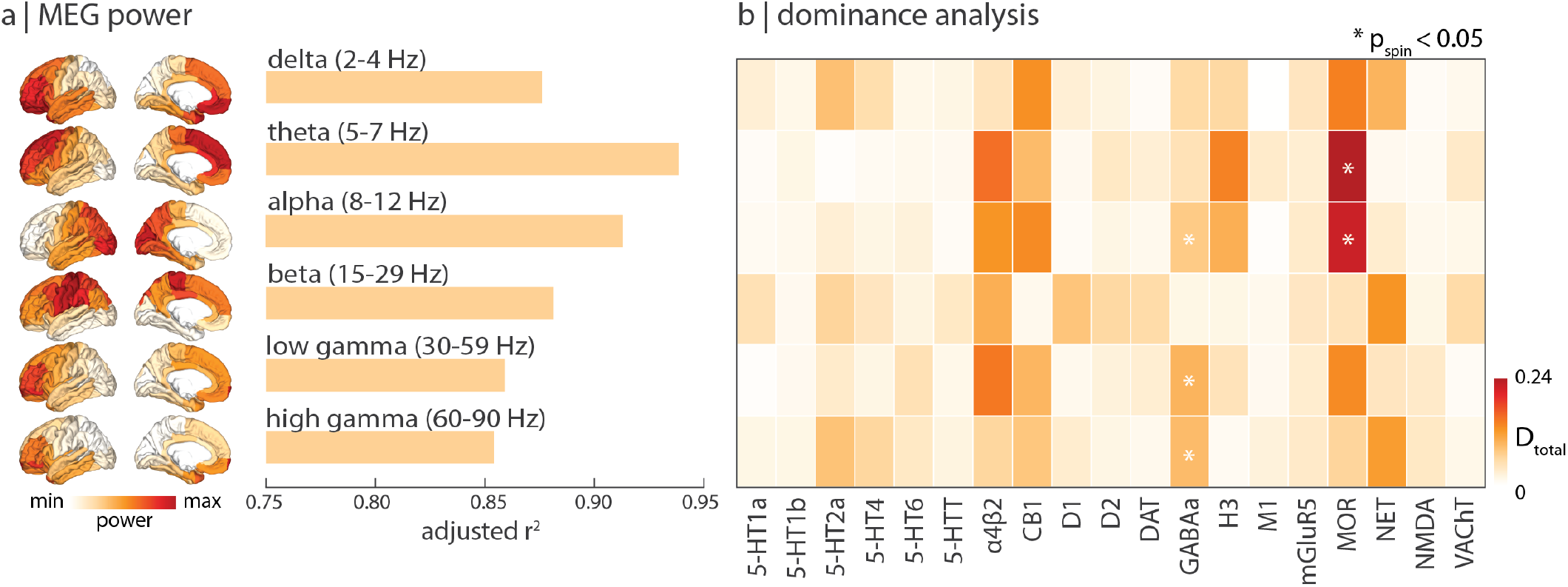
Receptor profiles shape oscillatory neural dynamics. We fit a multilinear regression model that predicts MEG-derived power distributions from receptor distributions. (a) Receptor distributions closely correspond to all six MEG-derived power bands (0.85 ≤ 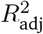 ≤ 0.94). (b) Dominance analysis reveals which receptors/transporters contribute most to the fit. Asterisks indicate significant dominance (*p*_spin_ < 0.05).

To determine which independent variables (receptors/transporters) contribute most to the fit, we applied dominance analysis, a technique that assigns a proportion of the final 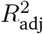 to each independent variable (Fig. 4b; [7]). Dominance was assessed against spin tests to identify neurotransmitter receptors that contribute to the fit above and beyond the effects of spatial autocorrelation. Notably, we find that the *μ*-opioid receptor (MOR) shows high dominance for theta and alpha frequency bands, consistent with previous reports [93, 95, 148, 150, 158]. Interestingly, the ionotropic GABA_A_ receptor is significantly dominant for frequency bands with fast timescales (alpha, low gamma, high gamma). The prominence of ionotropic receptors is also observed in the autoradiography dataset (see *Replication using autoradiography* and Fig. S3). Collectively, these results suggest that fast-acting ionotropic receptors promote neural oscillatory dynamics.

### Mapping receptors to cognitive function

Previously, we showed that receptor and transporter distributions follow the structural and functional organization of the brain, and that receptors are closely linked to neural dynamics. In this and the next subsections, we investigate how the spatial distribution of neurotransmitter receptors and transporters correspond to cognitive processes and disease vulnerability.

We used Neurosynth to derive meta-analytic task activation maps, which represent the probability that specific brain regions are activated during multiple cognitive tasks [151]. We selected a subset of 123 cognitive processes at the intersection of Neurosynth and the Cognitive Atlas [49, 96], and parcellated the data into the same 68-region atlas used for receptor maps, resulting in a region × cognitive process matrix of functional activations. We then applied partial least squares (PLS) analysis to identify a multivariate mapping between neurotransmitter receptors/transporters and functional activation maps (see *Methods* for details and Table S2 for the complete list of 123 cognitive terms; [64, 74]).

PLS analysis extracted a significant latent variable relating receptor/transporter densities to functional activation across the brain (*p*_spin_ = 0.036). The latent variable represents a spatial pattern of receptor distributions (receptor weights) and functional activations (cognitive weights) that together capture 44% of the covariance between the two datasets (Fig. 5a). Projecting the receptor density (functional activation) matrix back onto the receptor (cognitive) weights reflects how well a brain area exhibits the receptor and cognitive weighted pattern, which we refer to as “receptor scores” and “cognitive scores”, respectively (Fig. 5b, c). The receptor and cognitive score patterns reveal a sensory-fugal spatial gradient that reflects classes of laminar differentiation, separating limbic, paralimbic, and insular cortices from visual and somatosensory cortices (Fig. 5d; [76, 92]). We then cross-validated the correlation between receptor and cognitive scores using a distance-dependent method (mean out-of-sample *r* = 0.29; see *Methods* for details on the cross-validation). This result demonstrates a link between receptor distributions and cognitive specialization that is perhaps mediated by laminar differentiation and synaptic hierarchies.

**Figure 5.**
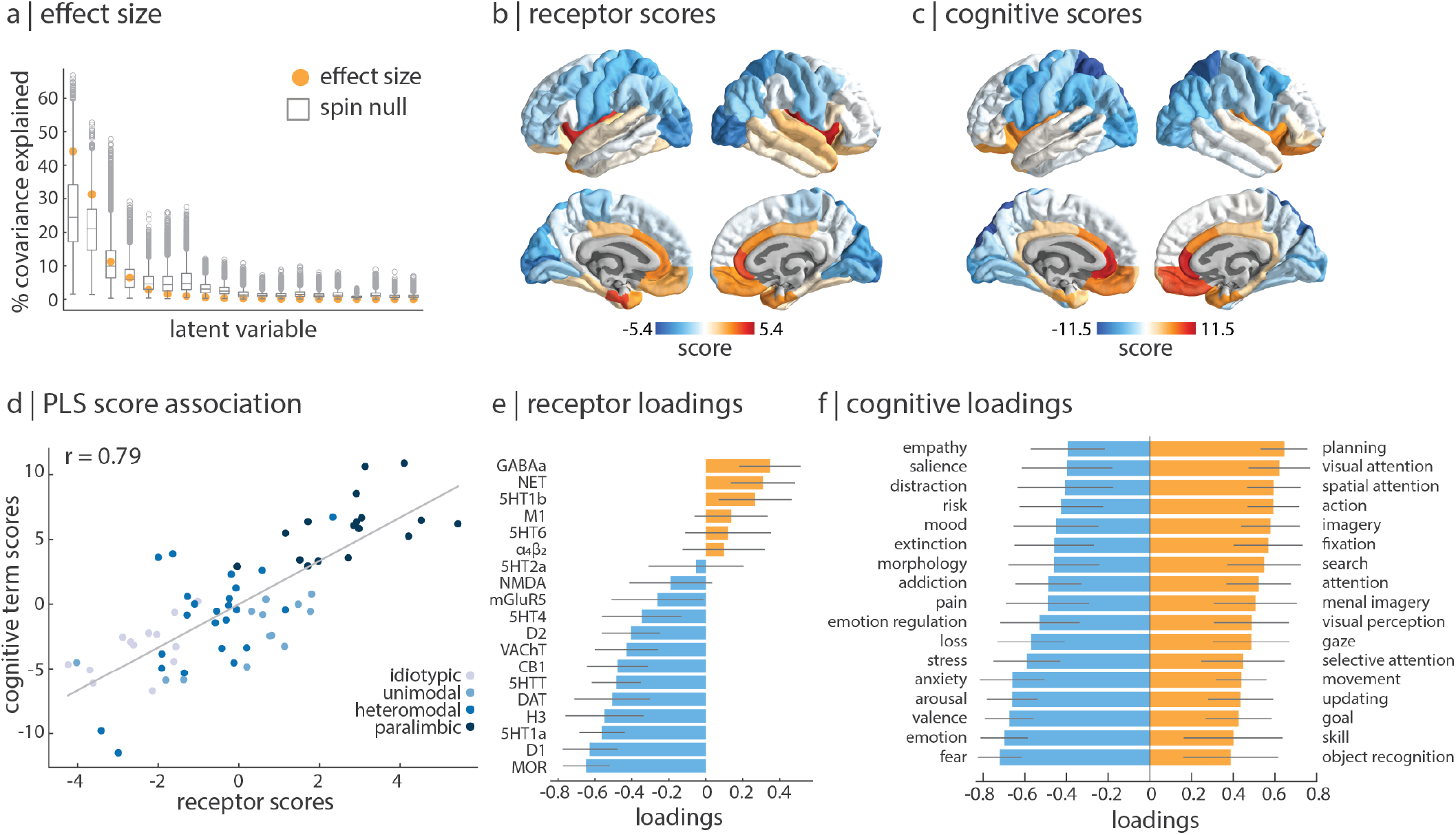
Mapping receptors to cognitive function. (a) Using partial least squares analysis (PLS), we find a significant latent variable that accounts for 44% of the covariation between receptor distributions and Neurosynth-derived cognitive functional activation (*p*_spin_ = 0.036, 10 000 repetitions). (b)–(c) This latent variable represents a pattern of coactivation between receptors (“receptor scores”) and cognitive terms (“cognitive scores”). (d) Receptor and cognitive scores are designed to correlate highly (*r* = 0.79, out-of-sample mean *r* = 0.32). Points are coloured according to their Mesulam classes of laminar differentiation, revealing a sensory-fugal gradient [76, 92]. (e) Receptor loadings are computed as the correlation between each receptor’s distribution across the cortex and the PLS-derived scores, and can be interpreted as the contribution of each receptor to the latent variable. (f) Similarly, cognitive loadings are computed as the correlation between each term’s functional activation across brain regions and the PLS-derived scores, and can be interpreted as the cognitive processes that contribute most to the latent variable. Here, only the 25% most positively and negatively loaded cognitive processes are shown. For all stable cognitive loadings, see Fig. S5 and for all 123 cognitive processes included in the analysis, see Table S2. 95% confidence intervals are estimated for receptor and cognitive loadings using bootstrap resampling (10 000 repetitions).

To identify the receptors and cognitive processes that contribute most to the spatial pattern in Fig. 5b and c, we correlated each variable with the score pattern (Fig. 5e – f; for all stable term loadings, see Fig. S5). This results in a “loading” for each receptor and cognitive process, where positively loaded receptors covary with positively loaded cognitive processes in positively scored brain regions, and vice versa for negative loadings. Interestingly, we find that the cognitive processes with the greatest negative loading are enriched for emotional and affective processes such as “fear”, “emotion”, and “valence”. Likewise, the neurotransmitter receptors with the greatest negative loading are broadly mood-related, including MOR (pain management), D_1_, and 5-HT_1A_ (mood regulation). On the other hand, cognitive processes with the greatest positive loading are related to sensory processing and interacting with the external environment, including “visual attention”, “spatial attention” and “action”. In other words, we find a patterning of receptor profiles that robustly separates areas involved in extrinsic function, including sensory-motor processing and attention, versus areas involved in affect and interoception. Collectively, these results demonstrate a direct link between cortex-wide molecular receptor distributions and functional specialization.

### Mapping receptors and transporters to disease vulnerability

Neurotransmitter receptors and transporters are implicated in multiple diseases and disorders. Identifying the neurotransmitter receptors/transporters that correspond to specific disorders is important for developing new therapeutic drugs. We therefore sought to relate neurotransmitter receptors and transporters to patterns of cortical thinning (a proxy for loss of neurons and synapses [146]) across a range of neurological, developmental, and psychiatric disorders. We used datasets from the ENIGMA consortium for a total of 13 disorders including: 22q11.2 deletion syndrome (22q) [131], attention deficit hyperactivity disorder (ADHD) [53], autism spectrum disorder (ASD) [142], idiopathic generalized epilepsy, right and left temporal lobe epilepsy [149], depression [114], obsessive-compulsive disorder (OCD) [15], schizophrenia [139], bipolar disorder (BD) [51], obesity [88], schizotypy [61], and Parkinson’s disease (PD) [65]. All cortical thinning maps were collected from adult patients, following identical processing protocols, for a total of over 21 000 scanned patients against almost 26 000 controls. We then fit a multiple regression model that predicts each disorder’s cortical thinning pattern from receptor and transporter distributions (Fig. 6), and evaluated each model using a distance-dependent cross-validation (Fig. S4; see *Methods* for details on the cross-validation).

**Figure 6.**
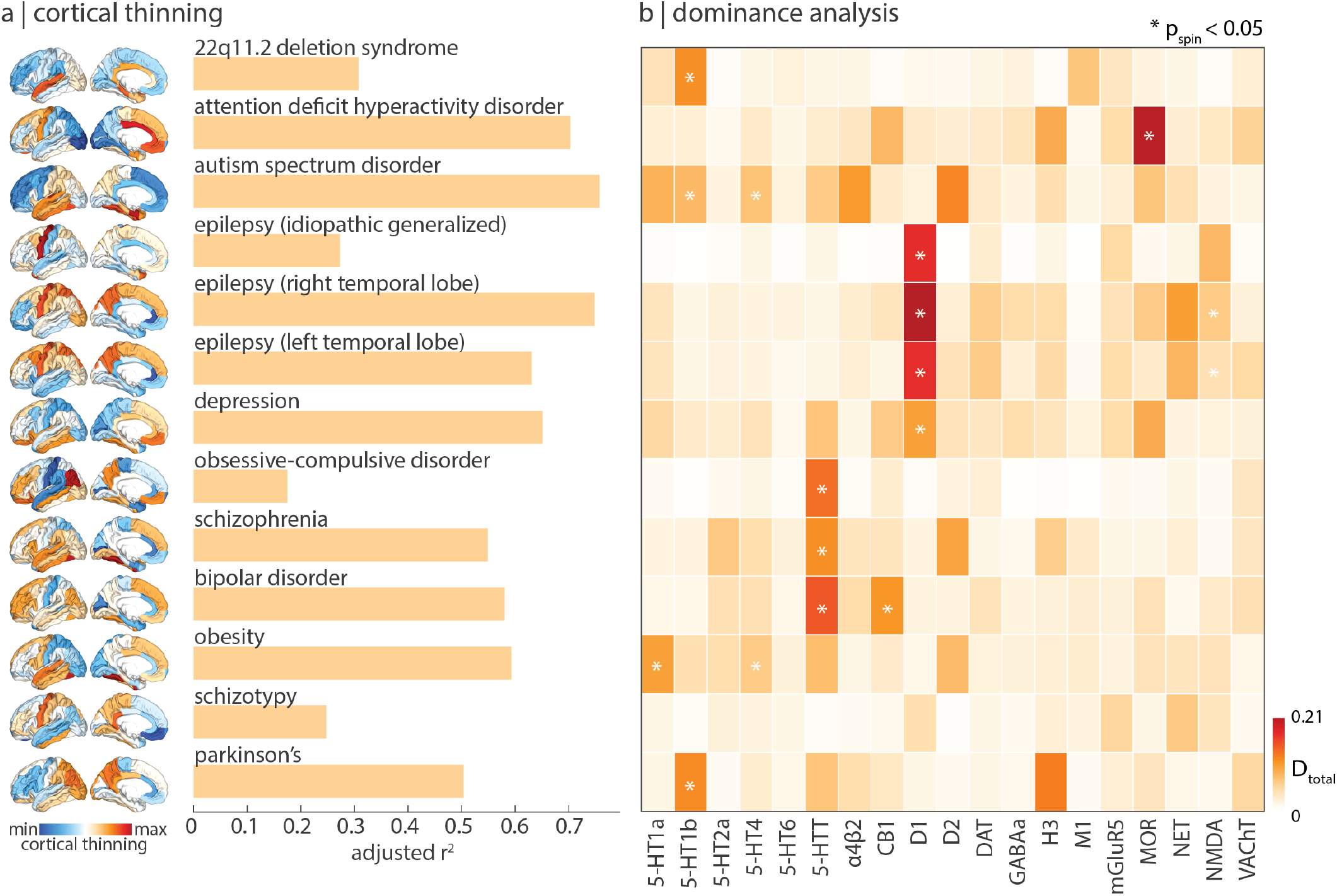
Mapping receptors to disease vulnerability. Using a multilinear model, neurotransmitter receptor/transporter distributions were fit to patterns of cortical thinning for thirteen neurological, psychiatric, and neurodevelopmental disorders, collected by the ENIGMA consortium [67, 134]. (a) Model fit varies: idiopathic generalized epilepsy, OCD, and schizotypy atrophy patterns show smaller fits, and ADHD, autism, and temporal lobe epilepsy atrophy patterns show greater fits. (b) Dominance analysis identified the neurotransmitter receptors and transporters whose spatial distribution most influence model fit, for each disorder separately. Asterisks indicate significant dominance (*p*_spin_ < 0.05).

Figure 6a shows how receptor distributions map onto cortical thinning patterns across multiple disorders. We find that some disorders are more heavily influenced by receptor distribution than others (0.17 < 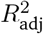 < 0.76). Idiopathic generalized epilepsy, OCD, and schizotypy show lower correspondence with receptor distributions, whereas ADHD, autism, and temporal lobe epilepsies show greater correspondence with receptor distributions. Interestingly, we find that serotonin transporter (5-HTT) distributions conform to cortical atrophy of psychiatric mood disorders such as OCD, schizophrenia, and bipolar disorder. Furthermore, MOR receptor distributions map onto cortical thinning patters of ADHD, consistent with findings from animal models [23, 105]. We also note that in some cases the analyses do not necessarily recover the expected relationships. For instance, in PD, the dopamine receptors are not implicated, likely because the analysis was restricted to cortex only. Additionally, 5-HTT is not significantly dominant towards depression, possibly because cortical thinning does not directly measure the primary pathophysiology associated with some brain diseases. Our results present an initial step towards a comprehensive “look-up table” that relates neurotransmitter systems to multiple brain disorders.

### Replication using autoradiography

In the present report, we comprehensively situate neurotransmitter receptor and transporter densities within the brain’s structural and functional architecture. However, estimates for neurotransmitter receptor densities are acquired from PET imaging alone, and the way in which densities are quantified varies across radioligands, image acquisition protocols, and preprocessing. Autoradiography is an alternative technique to measure receptor density, and captures local densities at a defined number of post-mortem brain sections [157]. Due to the high cost and labor intensity of acquiring autoradiographs, there does not yet exist a complete autoradiography 3-D cross-cortex atlas of receptors (but see [38]).

Nonetheless, we repeated the analyses in an autoradiography dataset of 15 neurotransmitter receptors across 44 cytoarchitectonically defined cortical areas, from three post-mortem brains [48, 155]. This set of 15 neurotransmitter receptors consists of a diverse set of ionotropic and metabotropic receptors, including excitatory glutamate, acetylcholine, and norepinephrine receptors (see Table S1 for a complete list of receptors and Fig. S6a for mean density of neurotransmitter receptors across the 44 regions). Correlations of receptor density distribution between every pair of receptors is shown in Fig. S6b, and receptor similarity is shown in Fig. S6c. Despite the alternate set of neurotransmitter receptors, we find that autoradiography-derived receptor similarity is significantly correlated with PET-derived receptor similarity (*r* = 0.32, *p* = 1.9 *×* 10^−14^; Fig. S6d). Additionally, we find that autoradiography-derived receptor densities follow similar architectural patterns as the PET-derived receptor densities. Within the autoradiography dataset, receptor similarity decays exponentially with distance and is significantly greater between structurally connected brain regions (*p* = 0.04), is non-significantly greater in regions within the same intrinsic network (*p*_spin_ = 0.11), and is significantly correlated with functional connectivity (*r* = 0.34, *p* = 9.4 × 10^−16^; Fig. S6e – f). As before, receptor information augments structure-function coupling in dorsolateral prefrontal and temporal regions (Fig. S6g).

Since the autoradiography dataset has a more diverse set of ionotropic and metabotropic receptors, we also asked whether we could replicate the dominance of ionotropic receptors for MEG oscillations. When we fit the fifteen autoradiography neurotransmitter receptors to MEG power, we find that AMPA, NMDA, GABA_A_, and *α*_4_*β*_2_—all ionotropic receptors—are most dominant, although only AMPA, NMDA, and GABA_A_ are significant (Fig. S3). This confirms that the fast oscillatory dynamics captured by MEG are closely related to the fluctuations in neural activity modulated by ionotropic neurotransmitter receptors.

Finally, we repeat analyses mapping receptor densities to cognitive functional activation and disease vulnerability. We find a similar topographic gradient linking autoradiography-derived receptor densities to Neurosynth-derived functional activations (Fig. S7a), as well as consistencies regarding the loadings of receptors (Fig. S7b) and cognitive processes (Fig. S7c). Next, when we map autoradiography-derived receptor densities to cortical thinning patterns of multiple disorders, we find prominent associations with receptors that were not included in the PET dataset, including a relationship between the ionotropic glutamate receptor kainate and the epilepsies (Fig. S8; [89]).

### Sensitivity and robustness analyses

Finally, to ensure results are not influenced by specific methodological choices, we repeated analyses using different parcellation resolutions, different receptor subsets, and we compared alternative PET tracers to the chosen PET tracers in the present report. Due to the low spatial resolution of PET tracer binding, we opted to present our main results using a coarse resolution of 68 Desikan-Killiany cortical regions. However, when using a parcellation resolution of 114 and 219 cortical regions [18], we find that the mean receptor density and receptor similarity remains consistent (Fig. S9). We next asked whether any single receptor or transporter disproportionately influences receptor similarity. To test this, we iteratively removed a single receptor/transporter from the dataset and recomputed the receptor similarity matrix. These 19 different receptor similarity matrices are all highly correlated with the original similarity matrix (*r >* 0.98), confirming that the correspondence between regional receptor profiles is not driven by a single neurotransmitter receptor/transporter.

Constructing a harmonized set of PET neurotransmitter receptor maps necessitated several methodological decisions. We combined PET maps from different research groups that used the same tracer, except for data from the Neurobiology Research Unit in Copenhagen (including [12] and [84]; see https://xtra.nru.dk/) because these images were converted from quantitative PET Units to units of density (pmol/mL) using autoradiography data. We combined two P943 (5HT1_B_ [30, 39, 113]), two FLB457 (D_2_ [109, 126]), three ABP688 (mGluR_5_ [28, 125]), and four FEOBV (VAChT [1, 10]) tracer maps separately, all of which were highly correlated within tracer groups (Fig. S1a). Next, when multiple tracers were available for the same receptor or transporter, we opted for maps constructed from a larger number of participants (Fig. S1b; see *Methods* for details).

Finally, we tested whether participant age affects the reported results. However, only mean age of individuals included in each tracer map was available. Therefore, we fit a linear model between the mean age of scanned participants contributing to each receptor/transporter tracer map and the z-scored receptor/transporter density, for each brain region separately. We then subtracted the relationship with age from the original receptor densities, resulting in an age-regressed receptor density matrix. We find that both age-regressed receptor density and age-regressed receptor similarity is highly correlated with the original receptor density/similarity (*r* = 0.81 and *r* = 0.98, respectively; Fig. S10), suggesting that age has negligible effect on the reported findings.

## Discussion

In the present report, we curate a comprehensive 3-D atlas of 19 neurotransmitter receptors and transporters. We demonstrate that chemoarchitecture is a key layer of the multi-scale organization of the brain. Neurotransmitter receptor profiles closely align with the structural connectivity of the brain and mediate its link with function, including neurophysiological oscillatory dynamics, and resting state hemodynamic functional connectivity. The overlapping topographic distributions of these receptors ultimately manifest as patterns of cognitive specialization and disease vulnerability.

A key question in neuroscience remains how the brain’s structural architecture gives rise to its function [6, 130]. The relationship between whole-brain structure and function has been viewed through the lens of “connectomics”, in which the brain’s structural or functional architectures are represented by regional nodes interconnected by structural and functional links. The key assumption of this model is that nodes are homogenous, effectively abstracting away important microarchitectural differences between regions. The present work is part of an emerging effort to annotate the connectome with molecular, cellular, and laminar attributes [145]. Indeed, recent work has incorporated microarray gene transcription [17, 49], cell types [4, 117], myelination [24, 25, 55], laminar differentiation [147], and intrinsic dynamics [43, 69, 79, 120] into structural and functional models of the brain.

Neurotransmitter receptors and transporters are an important molecular annotation for bridging brain structure to brain function. Neurotransmitter receptors support signal propagation across electrochemical synapses and tune neural gain [121, 123]. Despite their importance, a comprehensive cortical map of neurotransmitter receptors has remained elusive due to numerous methodological and data sharing challenges (but see the ongoing PET-BIDS effort as well as the OpenNeuro PET initiative at https://openneuropet.github.io/ [62, 86]). The present study is an ongoing Open Science grassroots effort to assemble harmonized high-resolution normative images of receptors and transporters that can be used to annotate connectomic models of the brain. This work builds on previous initiatives to map receptor densities using autoradiography, which has discovered prominent gradients of receptor expression in both human and macaque brains [36, 48, 155]. Importantly, we find consistent results between autoradiography and PET datasets, which is encouraging because the PET dataset consists of a different group of receptors and transporters, and has the added advantage of providing *in vivo* whole-brain data in large samples of healthy young participants.

We find that structurally connected areas have more similar receptor profiles, suggesting that neurotransmitter receptors are systematically aligned with network structure to regulate inter-regional communication. Indeed, we find a prominent link between receptor distribution and function, including correlated receptor similarity and functional connectivity, as well as greater receptor similarity within intrinsic functional networks. These results support the idea that the emergent functional architecture strongly depends on the underlying chemoarchitecture [122, 154]. Interestingly, we find that the canonical electrophysiological frequency bands can be captured by the overlapping topographies of multiple receptors, consistent with the notion that receptors influence function by tuning gain and synchrony between neuronal populations.

Since receptors modulate the link between structure and function, a natural next question is how receptor distributions relate to psychological processes. We find a prominent spatial gradient of receptor profiles that separates areas involved in extrinsic function, including sensory-motor processing and attention, versus areas involved in affect and interoception. This gradient maps on to the sensory-fugal synaptic hierarchy proposed by Mesulam [36, 75]. The concordance between the two maps is noteworthy because one is derived from the wiring patterns of the brain, while the other is derived from mapping receptors to task-based activations. This suggests that the brain is characterized by universal organizational principles that can be observed at multiple scales of description. Our results bridge the microscale and macroscale to reveal a molecular signature of psychological processes.

Finally, we discover a robust spatial concordance between multiple receptor maps and cortical thinning across a wide range of brain disorders. A key step toward developing therapies for specific syndromes is to reliably map them onto underlying neural systems. This goal is challenging because psychiatric and neurological nosology is built around clinical features, rather than neurobiological mechanisms [56]. Our results complement some previously established associations between disorders and neurotransmitter systems, and also reveal new associations. For instance, we find that the serotonin transporter is consistently implicated in OCD and bipolar disorder, consistent with the fact that selective serotonin reuptake inhibitors (SSRI) are frequently used to treat such disorders—although we do not find a significant spatial relationship between the serotonin transporter and depression. Additionally, we find that serotonin receptors are associated with obesity, consistent with the notion that serotonin systems regulate homeostatic and hedonic circuitry and are therefore implicated in food intake [141]. On the other hand, we find associations that have some preliminary support in the literature, but to our knowledge have not been conclusively established and adopted into clinical practice, including histamine H_3_ in Parkinson’s disease [102, 116], MOR in ADHD [23, 105], and D_1_ and NET in temporal lobe epilepsy [20, 44, 129]. Mapping disease phenotypes to receptor profiles will help to identify novel targets for pharmacotherapy [60]. This analysis is restricted to a single perspective of disease pathology (cortical thinning) and should be expanded in future work to encompass other forms of disease presentation as well as the effects of age and pathology on receptor/transporter density. The present work should be considered alongside some important methodological considerations. First, main analyses were conducted using PET images, which detect tracer uptake at a low spatial resolution and without laminar specificity. Although results were replicated using an autoradiography dataset, and in a finer parcellation resolution, a comprehensive atlas of laminar-resolved receptor density measurements is necessary to fully understand how regional variations in receptor densities affect brain structure and function [91]. Second, PET tracer maps were acquired around the world, in different participants, on different scanners, and using specific image acquisition and processing protocols recommended for each individual radioligand [85, 144]. To mitigate this challenge, we normalized the spatial distributions and focused only on analyses related to the relative spatial topographies of receptors as opposed to the absolute values. Third, the linear models used in the present analyses assume independence between observations and linear relationships between receptors [**?**]; we therefore employ spatial-autocorrelation preserving null models to account for the spatial dependencies between regions throughout the report. Fourth, analyses were conducted in the cortex only, due to data availability and well documented differences in tracer uptake between the cortex and subcortex [32, 136]. Altogether, a 3-D whole-brain comprehensive neurotransmitter receptor density dataset constructed using autoradiographs would be a valuable complement to the present work [38, 91, 155].

In summary, we assemble a normative 3-D atlas of neurotransmitter receptors in the human brain. We systematically map receptors to connectivity, dynamics, cognitive specialization, and disease vulnerability. Our work uncovers a fundamental organizational feature of the brain and provides new direction for a multi-scale systems-level understanding of brain structure and function.

## Methods

All code and data used to perform the analyses can be found at https://github.com/netneurolab/hansen_receptors. Volumetric PET images are included in neuromaps (https://github.com/netneurolab/neuromaps) where they can be easily converted between template spaces.

### PET data acquisition

Volumetric PET images were collected for 19 different neurotransmitter receptors and transporters across multiple studies. To protect patient confidentiality, individual participant maps were averaged within studies before being shared. Details of each study, the associated receptor/transporter, tracer, number of healthy participants, age, and reference with full methodological details can be found in Table 1. A more extensive table can be found in the supplementary material (Table S3) which additionally includes the PET camera, number of males and females, PET modelling method, reference region, scan length, modelling notes, and additional references, if applicable. Note that three tracer maps were shared prior to publication, but contact information is available for the corresponding authors (Table S3). In all cases, only healthy participants were scanned. Images were acquired using best practice imaging protocols recommended for each radioligand [85]. Altogether, the images are an estimate proportional to receptor densities and we therefore refer to the measured value (i.e. binding potential, tracer distribution volume) simply as density. Note that the NMDA receptor tracer ([^18^F]GE-179) binds to open (i.e. active) NMDA receptors [73, 115]. PET images were all registered to the MNI-ICBM 152 nonlinear 2009 (version c, asymmetric) template, then parcellated to 68, 114, and 219 regions according to the Desikan-Killiany atlas [18, 26]. Receptors and transporters with more than one mean image of the same tracer (i.e. 5-HT_1B_, D_2_, mGluR_5_, and VAChT) were combined using a weighted average. Finally, each tracer map corresponding to each receptor/transporter was z-scored across regions and concatenated into a final region by receptor matrix of relative densities.

In some cases, more than one tracer map was available for the same neurotransmitter receptor/transporter. We show the comparisons between tracers in Fig. S1b for the following neurotransmitter receptors/transporters: 5-HT1_A_ [12, 113], 5-HT1_B_ [12, 39, 113], 5-HT2_A_ [12, 113, 133], 5-HTT [12, 113], CB_1_ [68, 87], D_2_ [2, 57, 109, 126], DAT [29, 112], GABA_A_ [29, 84], MOR [59, 135], and NET [27, 50]. Here we make some specific notes: (1) 5-HTT and GABA_A_ involve comparisons between the same tracers (DASB and flumazenil, respectively) but one map is converted to density using autoradiography data (see [12] and [84]) and the other is not [29, 30, 113]; (2) raclopride is a popular D_2_ tracer but has unreliable binding in the cortex, and is therefore an inappropriate tracer to use for mapping D_2_ densities in the cortex, but we show its comparison to FLB457 and another D_2_ tracer, fallypride, for completeness [2, 22, 57]; (3) the chosen carfentanil (MOR) map was collated across carfentanil images in the PET Turku Centre database—since our alternative map is a partly overlapping subset of participants, we did not combine the tracers into a single mean map [59, 135].

### Autoradiography receptor data acquisition

Receptor autoradiography data were originally acquired as described in [155]. 15 neurotransmitter receptor densities across 44 cytoarchitectonically defined areas were collected in three post-mortem brains (age range: 72–77, 2 males). See Table S1 for a complete list of receptors included in the autoradiography dataset, Supplementary Table 2 in [155] for the originally reported receptor densities, and https://github.com/AlGoulas/receptor_principles for machine-readable Python *numpy* files of receptor densities [48]. To best compare PET data analyses with the autoradiography dataset, a region-to-region mapping was manually created between the 44 available cortical areas in the autoradiography dataset and the 34 left hemisphere cortical Desikan Killiany regions. In only one case (the insula) was there no suitable mapping between the autoradiography data and the Desikan Killiany atlas. As such, the 44-region autoradiography atlas was converted to 33 Desikan Killiany left hemisphere regions (all but the insula). Finally, receptor densities were z-scored and averaged across supragranular, granular, and infragranular layers, to create a single map of receptor densities across the cortex.

### Structural and functional data acquisition

Structural and functional data were collected at the Department of Radiology, University Hospital Center and University of Lausanne, on *n* = 70 healthy young adults (27 females, 28.8 ± 9.1 years). Informed consent was obtained from all participants and the protocol was approved by the Ethics Committee of Clinical Research of the Faculty of Biology and Medicine, University of Lausanne. The scans were performed in a 3-T MRI scanner (Trio; Siemens Medical), using a 32-channel head coil. The protocol included (1) a magnetization-prepared rapid acquisition gradient echo (MPRAGE) sequence sensitive to white/grey matter contrast (1 mm in-plane resolution, 1.2 mm slice thickness), (2) a DSI sequence (128 diffusion-weighted volumes and a single b0 volume, maximum b-value 8 000s/mm^2^, 2.2 × 2.2 × 3.0 mm voxel size), and (3) a gradient echo-planar imaging (EPI) sequence sensitive to blood-oxygen-level-dependent (BOLD) contrast (3.3 mm in-plane resolution and slice thickness with a 0.3 mm gap, TR 1 920 ms, resulting in 280 images per participant). Participants were not subject to any overt task demands during the fMRI scan. The Lausanne dataset is available at https://zenodo.org/record/2872624#.XOJqE99fhmM.

### Structural network reconstruction

Grey matter was parcellated according to the 68-region Desikan-Killiany cortical atlas [26]. Structural connectivity was estimated for individual participants using deterministic streamline tractography. The procedure was implemented in the Connectome Mapping Toolkit [21], initiating 32 streamline propagations per diffusion direction for each white matter voxel. A group-consensus binary network was constructed using a method that preserves the density and edge-length distributions of the individual connectomes [14, 77, 78]. The density for the final 68-region structural connectome was 24.6%.

### Functional network reconstruction

Functional MRI data were preprocessed using procedures designed to facilitate subsequent network exploration [97]. fMRI volumes were corrected for physiological variables, including regression of white matter, cerebrospinal fluid, and motion (3 translations and 3 rotations, estimated by rigid body coregistration). BOLD time series were then subjected to a low-pass filter (temporal Gaussian filter with full width at half maximum equal to 1.92 s). The first four time points were excluded from subsequent analysis to allow the time series to stabilize. Motion “scrubbing” was performed as described by [97]. The data were parcellated according to the same 68-region Desikan-Killiany atlas used for the structural network. Individual functional connectivity matrices were defined as zero-lag Pearson correlation among the fMRI BOLD time series. A group-consensus functional connectivity matrix was estimated as the mean connectivity of pairwise connections across individuals. Note that one individual did not undergo an fMRI scan and therefore the functional connectome was composed of *n* = 69 participants.

### Structure-function coupling

Structure-function coupling was computed as per [143]. At each brain region, mutlilinear regression model was used to predict functional connectivity from three measures of the binary structural connectome. These structural measures were the Euclidean distance, shortest path length, and communicability between the region of interest and every other region. In the receptor-informed model, receptor similarity between the region of interest and every other region was included as an additional independent variable. Path length was computed using the Python package *bctpy* and is defined as the shortest contiguous set of edges between two brain regions, which represents a form of routing communication [104]. Communicability is defined as the weighted average of all walks and paths between two brain regions, and represents diffusive communication [33]. Coupling was defined as the adjusted *R*^2^ of the model.

### MEG power

6-minute resting state eyes-open magenetoen-cephalography (MEG) time-series were acquired from the Human Connectome Project (HCP, S1200 release) for 33 unrelated participants (age range 22—35, 17 males) [45, 140]. Complete MEG acquisition protocols can be found in the HCP S1200 Release Manual. For each participant, we computed the power spectrum at the vertex level across six different frequency bands: delta (2–4 Hz), theta (5–7 Hz), alpha (8–12 Hz), beta (15–29 Hz), low gamma (30–59 Hz), and high gamma (60–90 Hz), using the open-source software, Brainstorm [132]. The preprocessing was performed by applying notch filters at 60, 120, 180, 240, and 300 Hz, and was followed by a high-pass filter at 0.3 Hz to remove slow-wave and DC-offset artifacts. Preprocessed sensor-level data was used to obtain a source estimation on HCP’s fsLR4k cortex surface for each participant. Head models were computed using overlapping spheres and the data and noise covariance matrices were estimated from the resting state MEG and noise recordings. Brainstorm’s linearly constrained minimum variance (LCMV) beamformers method was applied to obtain the source activity for each participant. Welch’s method was then applied to estimate power spectrum density (PSD) for the source-level data, using overlapping windows of length 4 seconds with 50% overlap. Average power at each frequency band was then calculated for each vertex (i.e. source). Source-level power data was then parcellated into 68, 114, and 219 cortical regions for each frequency band [18].

### ENIGMA cortical thinning maps

The ENIGMA (Enhancing Neuroimaging Genetics through Meta-Analysis) Consortium is a data-sharing initiative that relies on standardized image acquisition and processing pipelines, such that disorder maps are comparable [134]. Patterns of cortical thinning were collected for thirteen neurological, neurodevelopmental, and psychiatric disorders from the ENIGMA consortium and the Enigma toolbox (https://github.com/MICA-MNI/ENIGMA; [66]) including: 22q11.2 deletion syndrome (22q) [131], attention deficit hyperactivity disorder (ADHD) [53], autism spectrum disorder (ASD) [142], idiopathic generalized epilepsy [149], right temporal lobe epilepsy [149], left temporal lobe epilepsy [149], depression [114], obsessive-compulsive disorder (OCD) [15], schizophrenia [139], bipolar disorder (BD) [51], obesity [88], schizotypy [61], and Parkinson’s disease (PD) [65]. Altogether, over 21 000 patients were scanned across the thirteen disorders, against almost 26 000 controls. The values for each map are z-scored effect sizes (Cohen’s *d*) of cortical thickness in patient populations versus healthy controls. For visualization purposes, data are inverted such that larger values represent greater cortical thinning. Imaging and processing protocols can be found at http://enigma.ini.usc.edu/protocols/.

### Dominance analysis

Dominance analysis seeks to determine the relative contribution (“dominance”) of each independent variable to the overall fit (adjusted *R*^2^) of the multiple linear regression model (https://github.com/dominance-analysis/dominance-analysis [7, 16]). This is done by fitting the same regression model on every combination of input variables (2^*p*^ − 1 submodels for a model with *p* input variables). Total dominance is defined as the average of the relative increase in *R*^2^ when adding a single input variable of interest to a submodel, across all 2^*p*^ − 1 submodels. The sum of the dominance of all input variables is equal to the total adjusted *R*^2^ of the complete model, making total dominance an intuitive measure of contribution. Significant dominance was assessed using the spin test (see *Null models*), whereby dominance analysis was repeated between a spun dependent variable and the original independent variables (1 000 repetitions).

### Distance-dependent cross-validation

The robustness of each multilinear model was assessed by cross-validating the model by using a distance-dependent method [49]. For each brain region, we select the 75% closest regions as the training set, and the remaining 25% of brain regions as the test set, for a total of 68 repetitions. This stratification procedure minimizes the dependence among the two sets due to spatial autocorrelation. The model was fit on the training set, and the predicted test-set output variable (either MEG power or cortical thinning) was correlated to the empirical test set values. The distribution of Pearson’s correlations between predicted and empirical power/cortical thinning across all repetitions (i.e. all brain regions) can be found in Fig. S2 and Fig. S4. Cross-validation of the partial least squares model is similarly conducted; more details can be found under *Partial least squares analysis*.

### Cognitive meta-analytic activation

Probabilistic measures of the association between voxels and cognitive processes were obtained from Neurosynth, a meta-analytic tool that synthesizes results from more than 15 000 published fMRI studies by searching for high-frequency key words (such as “pain” and “attention”) that are published alongside fMRI voxel coordinates (https://github.com/neurosynth/neurosynth, using the volumetric association test maps [151]). This measure of association is the probability that a given cognitive process is reported in the study if there is activation observed at a given voxel. Although more than a thousand cognitive processes are reported in Neurosynth, we focus primarily on cognitive function and therefore limit the terms of interest to cognitive and behavioural terms. These terms were selected from the Cognitive Atlas, a public ontology of cognitive science [96], which includes a comprehensive list of neurocognitive processes and has been previously used in conjunction with Neurosynth [3]. We used 123 terms, ranging from umbrella terms (“attention”, “emotion”) to specific cognitive processes (“visual attention”, “episodic memory”), behaviours (“eating”, “sleep”), and emotional states (“fear”, “anxiety”). The coordinates reported by Neurosynth were parcellated according to the 68-node Desikan Killiany atlas and z-scored. The probabilistic measure reported by Neurosynth can be interpreted as a quantitative representation of how regional fluctuations in activity are related to psychological processes. The full list of cognitive processes is shown in Table S2.

### Partial least squares analysis

Partial least squares analysis (PLS) was used to relate neurotransmitter receptor distributions to functional activation. PLS is an unsupervised multivariate statistical technique that decomposes the two datasets into orthogonal sets of latent variables with maximum covariance [64, 74]. The latent variables consist of receptor weights, cognitive weights, and a singular value which represents the covariance between receptor distributions and functional activations that is explained by the latent variable. Receptor and cognitive scores are computed by projecting the original data onto the respective weights, such that each brain region is assigned a receptor and cognitive score. Finally, receptor loadings are computed as the Pearson’s correlation between receptor densities and receptor scores, and vice versa for cognitive loadings.

The significance of the latent variable was assessed on the singular value, against the spin-test (see *Null models*). In the present report, only the first latent variable was significant; remaining latent variables were not analyzed further. Finally, the correlation between receptor and cognitive scores was cross-validated (see *Distance-dependent cross-validation*). After fitting PLS on the training set and correlating the ensuing receptor and cognitive scores, the test set was projected onto the training set-derived weights and the test set scores were correlated. The empirical correlation between receptor and cognitive scores across all brain regions was *r* = 0.79, the mean training set correlation was 0.86, and the mean test set correlation was 0.29.

### Null models

Spatial autocorrelation-preserving permutation tests were used to assess statistical significance of associations across brain regions, termed “spin tests” [3, 70]. We created a surface-based representation of the parcellation on the FreeSurfer fsaverage surface, via files from the Connectome Mapper toolkit (https://github.com/LTS5/cmp). We used the spherical projection of the fsaverage surface to define spatial coordinates for each parcel by selecting the coordinates of the vertex closest to the center of the mass of each parcel [143]. These parcel coordinates were then randomly rotated, and original parcels were reassigned the value of the closest rotated parcel (10 000 repetitions). Parcels for which the medial wall was closest were assigned the value of the next most proximal parcel instead. The procedure was performed at the parcel resolution rather than the vertex resolution to avoid upsampling the data, and to each hemisphere separately.

A second null model was used to test whether receptor similarity is greater in connected regions than unconnected regions. This model generates a null structural connectome that preserves the density, edge length, and degree distributions of the empirical structural connectome [13, 46, 103]. Briefly, edges were binned according to Euclidean distance. Within each bin, pairs of edges were selected at random and swapped. This procedure was then repeated 10 000 times. To compute a *p*-value, the mean receptor similarity of unconnected edges was subtracted from the mean receptor similarity of connected edges, and this difference was compared to a null distribution of differences computed on the rewired networks.

## Acknowledgment

We thank Vincent Bazinet, Zhen-Qi Liu, Filip Milisav, Laura Suarez, Bertha Vazquez-Rodriguez, and Mingze Li for their comments and suggestions on the manuscript. This research was undertaken thanks in part to funding from the Canada First Research Excellence Fund, awarded to McGill University for the Healthy Brains for Healthy Lives initiative. BM acknowledges support from the Natural Sciences and Engineering Research Council of Canada (NSERC Discovery Grant RGPIN #017-04265) and from the Canada Research Chairs Program. JYH acknowledges support from the Helmholtz International BigBrain Analytics & Learning Laboratory, the Natural Sciences and Engineering Research Council of Canada, and the Fonds de reserches de Québec. The funders had no role in study design, data collection and analysis, decision to publish or preparation of the manuscript.

---

**Figure S1.**
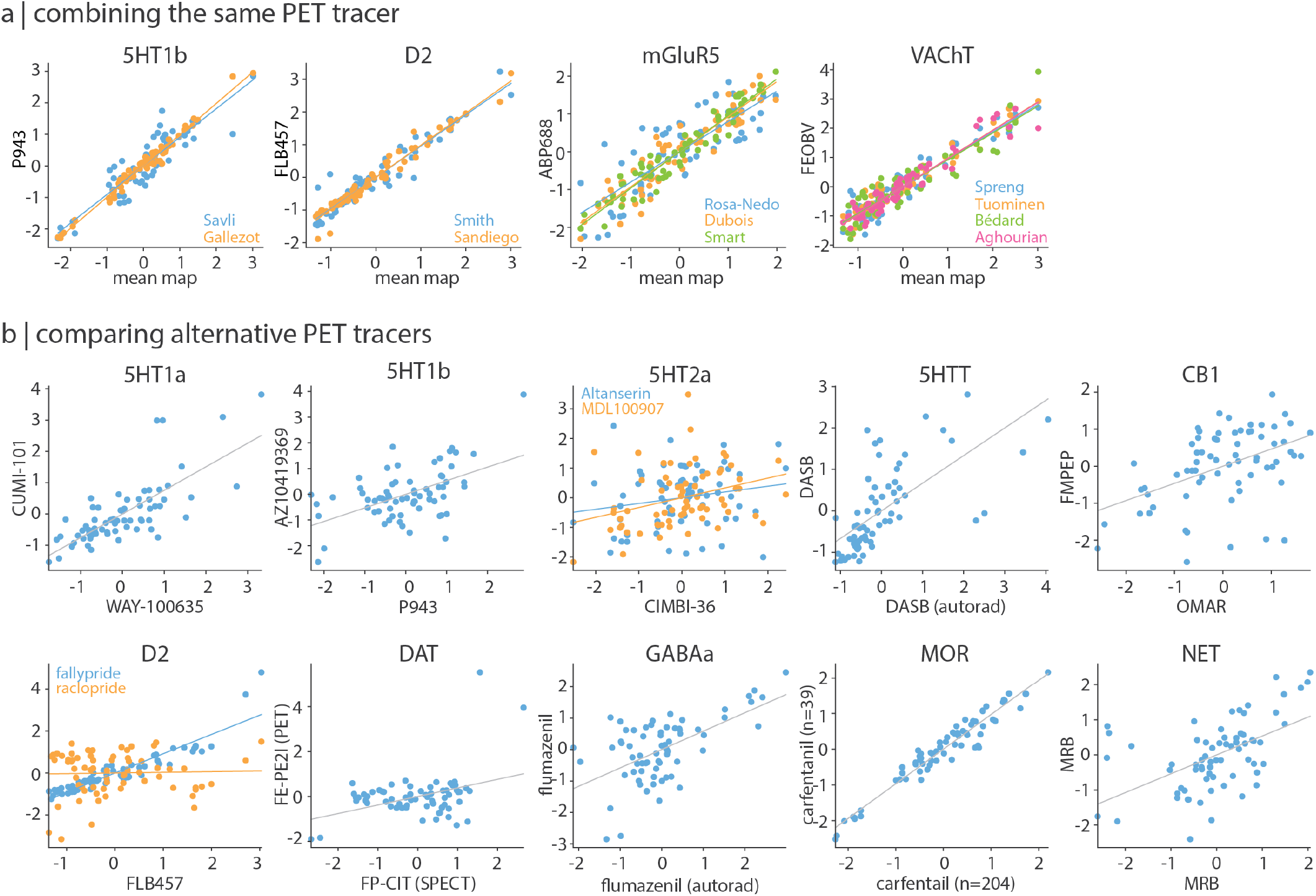
Comparing different PET tracer images. (a) PET maps of the same tracer were combined into a single average receptor/transporter map. Each individual PET tracer map (*y*-axis) is highly correlated to the mean map (*x*-axis). Names indicate the source of each PET map; see Table 1. (b) Multiple PET tracers were available for certain receptors/transporters. Scatter plots show the correlation between the selected tracer map (*x*-axis) and alternative maps (*y*-axis).

**Figure S2.**
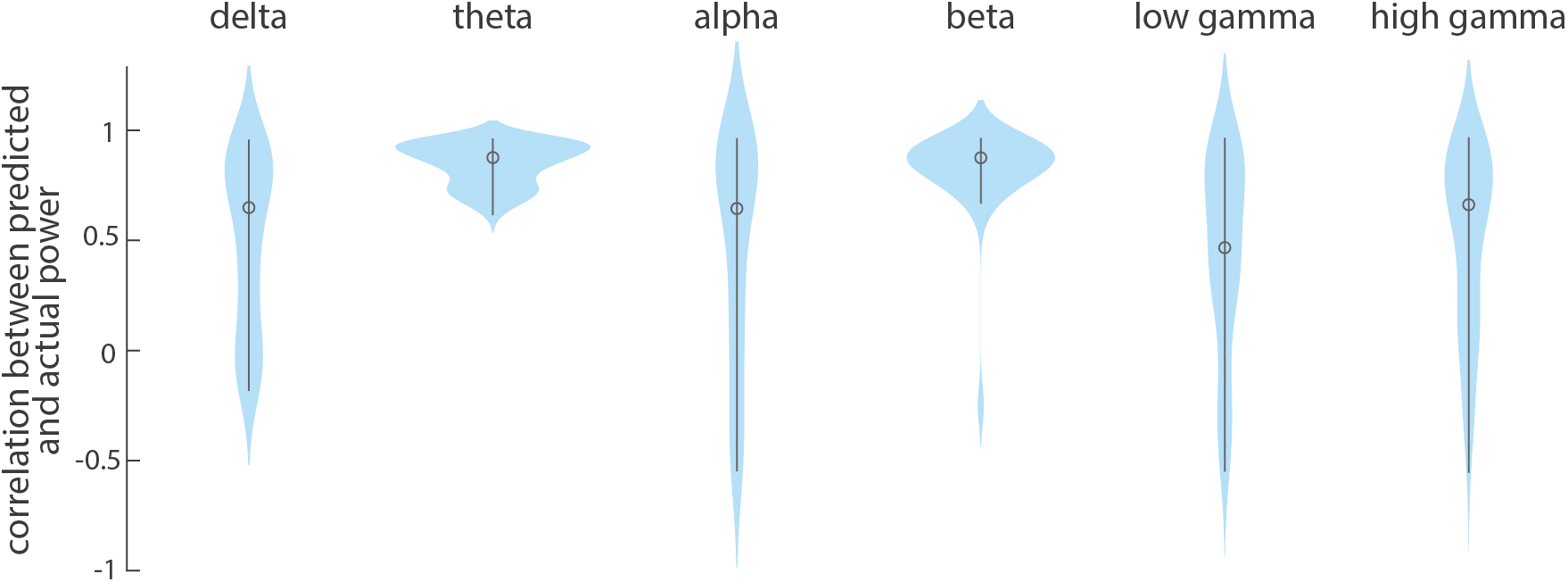
Cross-validating models that predict MEG power distribution from receptor/transporter densities. All six multilinear models between receptor/transporter densities and MEG power distributions were cross-validated using a distance-dependent method. This method selects the 25% of regions closest to a source-region as a training set and the remaining 75% of regions as the test set. The procedure is repeated for each brain region as the source region (68 iterations). We assessed the prediction by correlating predicted power to the empirical power in the test set.

**Figure S3.**
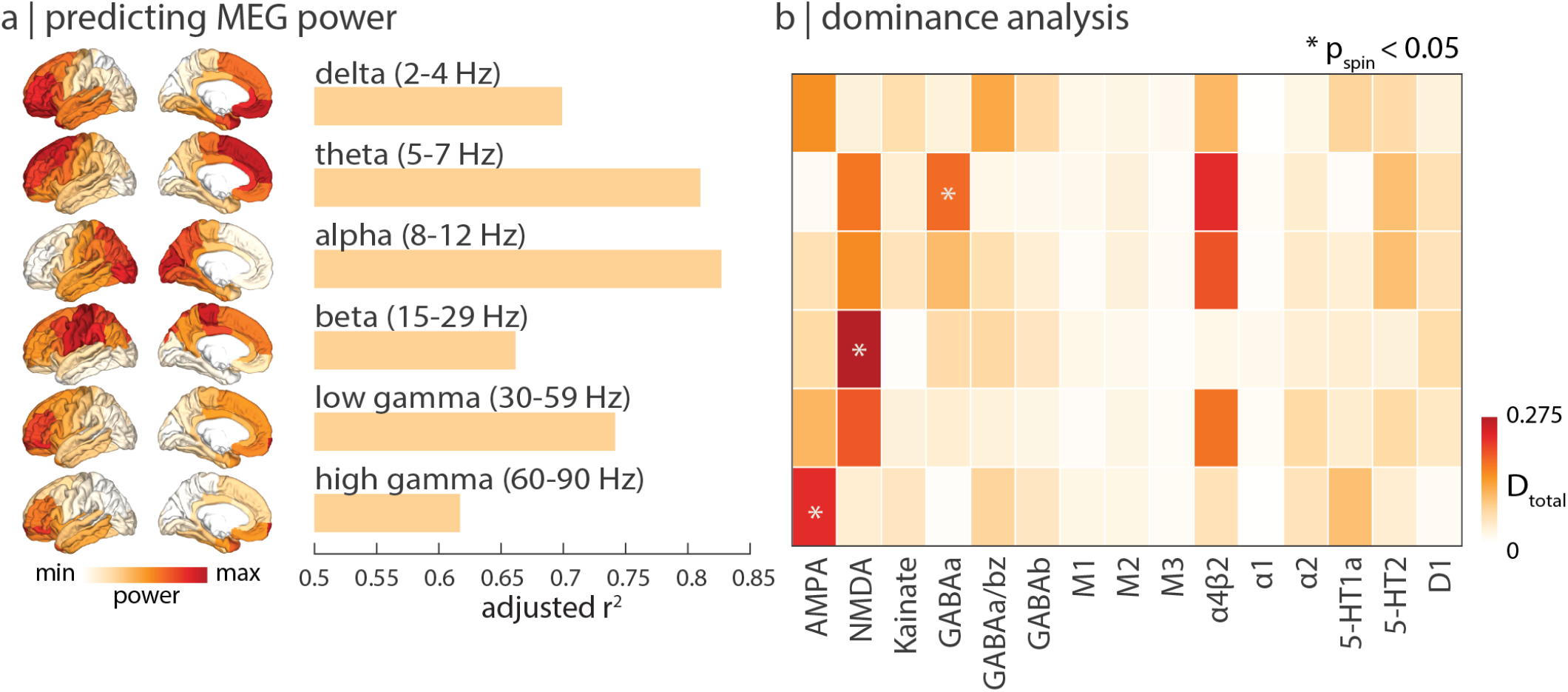
Excitatory ionotropic receptor densities shape neural dynamics. Multilinear regression models were fit between autoradiography-derived neurotransmitter receptor densities and MEG power, done analogously in Fig. 4. (a) Autoradiography-derived receptor densities map closely to neural dynamics. (b) Dominance analysis was applied to distribute the overall fit (adjusted *R*^2^) across the independent variables (receptors), revealing that excitatory ionotropic receptors contribute most to neural dynamics. Asterisks indicate significant dominance *p*_spin_ < 0.05.

**Figure S4.**
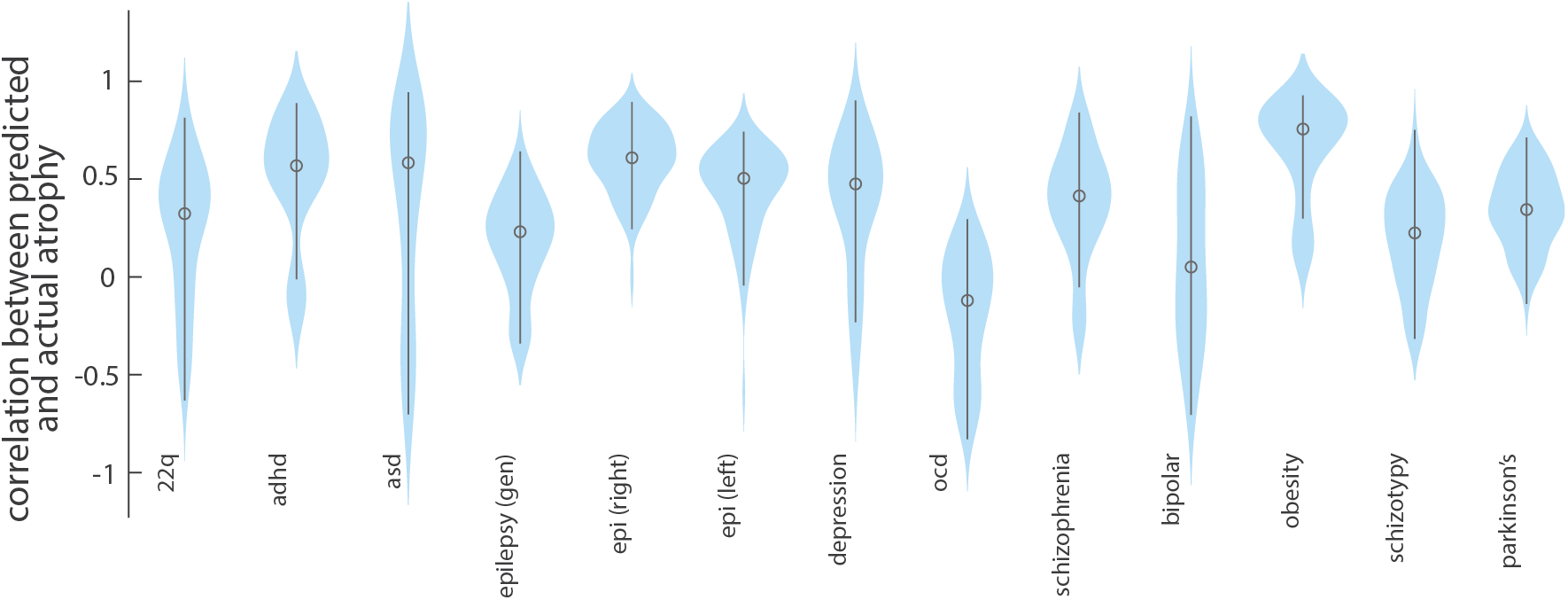
Cross-validating models that predict disorder-specific cortical thinning from receptor/transporter densities. All thirteen multilinear models between receptor/transporter densities and disorder-specific cortical thinning were cross-validated using a distance-dependent method. This method selects the 25% of regions closest to a source-region as a training set and the remaining 75% of regions as the test set. The procedure is repeated for each brain region as the source region (68 iterations). We assessed the prediction by correlating predicted atrophy to the empirical atrophy in the test set.

**Figure S5.**
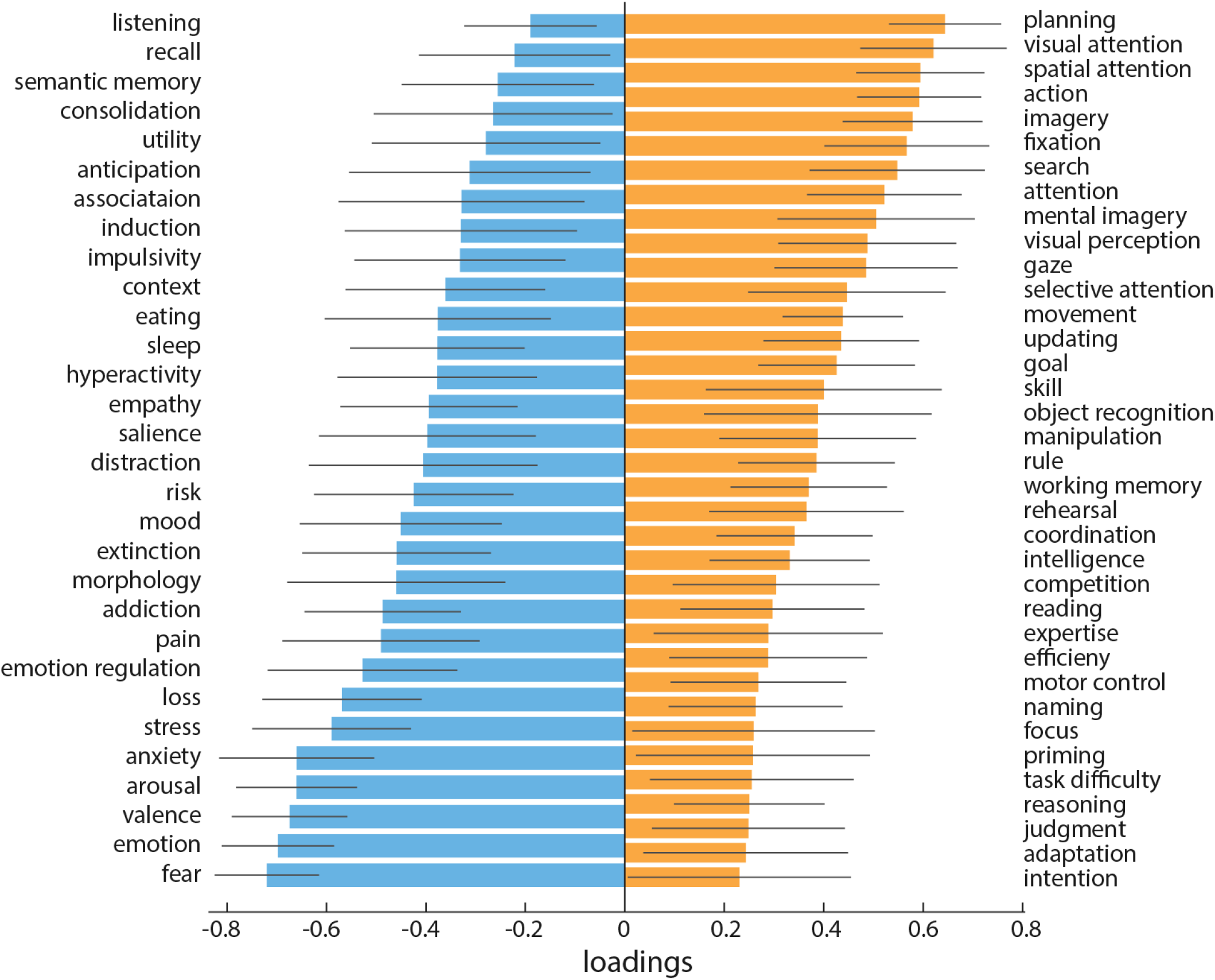
Neurosynth cognitive loadings. The loading for each cognitive process is calculated as the Pearson’s correlation between functional activations across brain regions and PLS-derived receptor scores. Error bars indicate bootstrap-estimated 95% confidence intervals (10 000 bootstrap samples). All cognitive processes with a confidence interval that changes sign are excluded.

**Figure S6.**
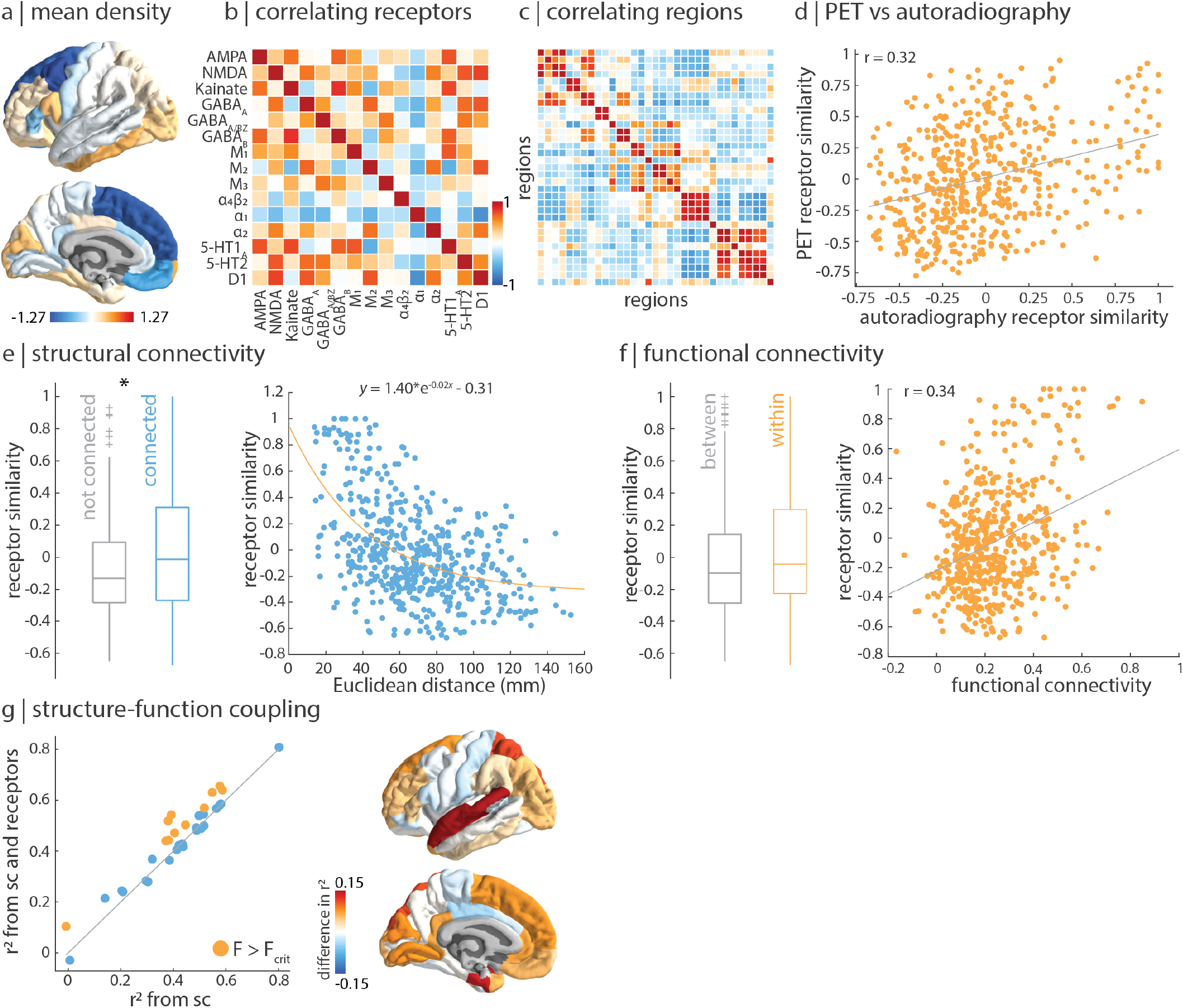
Autoradiography-informed neurotransmitter receptor densities follow similar organizational principles as PET-informed neurotransmitter receptor densities. Autoradiography images of fifteen neurotransmitter receptors across three postmortem brains were acquired by [155]. (a) Mean z-scored density, excluding the insula for which no data was available. (b) Pearson’s correlation between the receptor distribution across *n* = 33 brain regions for every pair of receptors. (c) The receptor similarity matrix is constructed by correlating receptor fingerprints at each pair of brain regions. (d) PET-derived receptor similarity is correlated to autoradiography-derived receptor similarity (*r* = 0.32, *p* = 1.9 × 10^−14^). (e) Receptor similarity is significantly greater between pairs of regions that are physically connected, against a degree- and edge-length-preserving null model (left; *p* = 0.04 [13]), and decays exponentially with Euclidean distance (right). (f) Receptor similarity is non-significantly greater in regions within the same functional network as opposed to between functional networks (left; *p*_spin_ = 0.11), and is correlated to functional connectivity (right; *r* = 0.34, *p* = 9.4 × 10^−16^). (g) Consistent with PET-derived results, receptor similarity augments structure-function coupling in temporal and prefrontal regions. Asterisks in panel (e) denote significance. Boxplots in (e) and (f) represent the 1st, 2nd (median) and 3rd quartiles, whiskers represent the non-outlier end-points of the distribution, and crosses represent outliers.

**Figure S7.**
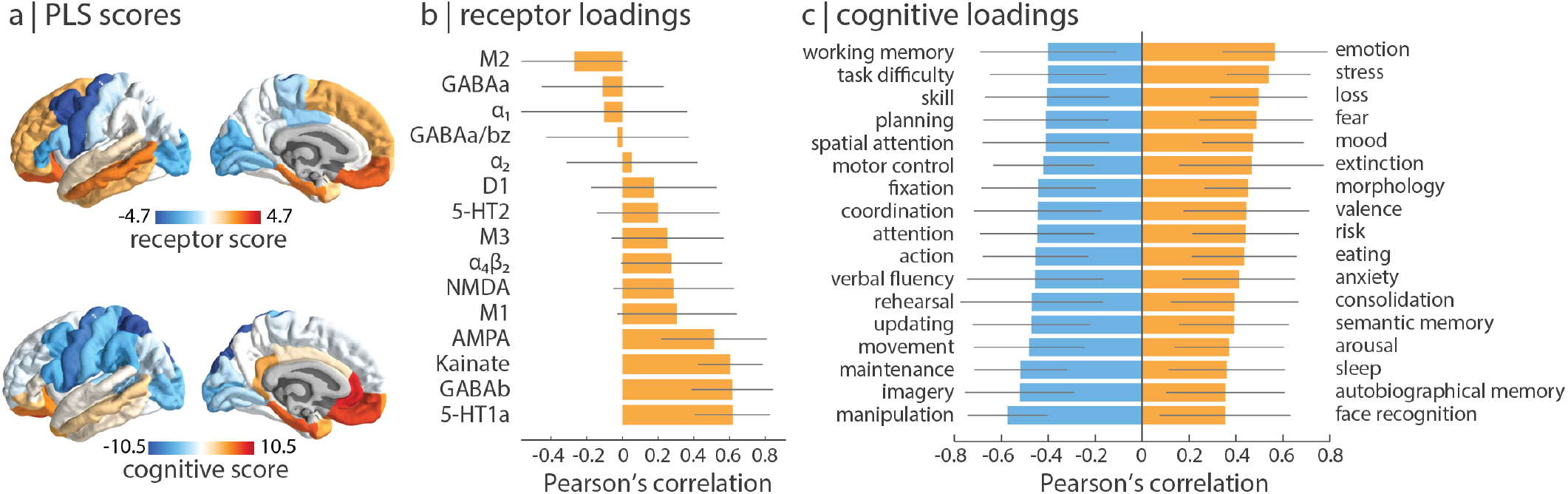
Mapping autoradiography-derived receptors to cognition. Partial least squares analysis was applied to autoradiography-derived receptor densities and Neurosynth-derived cognitive functional activations, done analogously in Fig. 5. (a) Receptor (top) and cognitive (bottom) score patterns follow a similar sensory-fugal gradient. (b) Receptor loadings are defined as the Pearson’s correlation between each receptor’s distribution across the cortex and the PLS-derived receptor scores and can be interpreted as the contribution of each receptor to the latent variable. (c) Cognitive loadings are shown for most positively- and negatively-loaded cognitive processes. 95% confidence intervals are estimated for receptor and cognitive loadings using bootstrap resampling (10 000 repetitions).

**TABLE S1.** Neurotransmitter receptors included in the autoradiography dataset.

| Receptor | Neurotransmitter | Excitatory/Inhibitory | Ionotropic/Metabotropic |
| --- | --- | --- | --- |
| AMPA | glutamate | excitatory | ionotropic |
| NMDA | glutamate | excitatory | ionotropic |
| Kainate | glutamate | excitatory | ionotropic |
| GABA<sub>A</sub> | GABA | inhibitory | ionotropic |
| GABA<sub>A/BZ</sub> | GABA | inhibitory | ionotropic |
| GABA<sub>B</sub> | GABA | inhibitory | metabotropic |
| M<sub>1</sub> | acetylcholine | excitatory | metabotropic |
| M<sub>2</sub> | acetylcholine | inhibitory | metabotropic |
| M<sub>3</sub> | acetylcholine | excitatory | metabotropic |
| $\alpha_4\beta_2$ | acetylcholine | excitatory | ionotropic |
| $\alpha_1$ | norepinephrine | excitatory | metabotropic |
| $\alpha_2$ | norepinephrine | inhibitory | metabotropic |
| 5-HT<sub>1A</sub> | serotonin | inhibitory | metabotropic |
| 5-HT<sub>2</sub> | serotonin | excitatory | metabotropic |
| D<sub>1</sub> | dopamine | excitatory | metabotropic |

**Figure S8.**
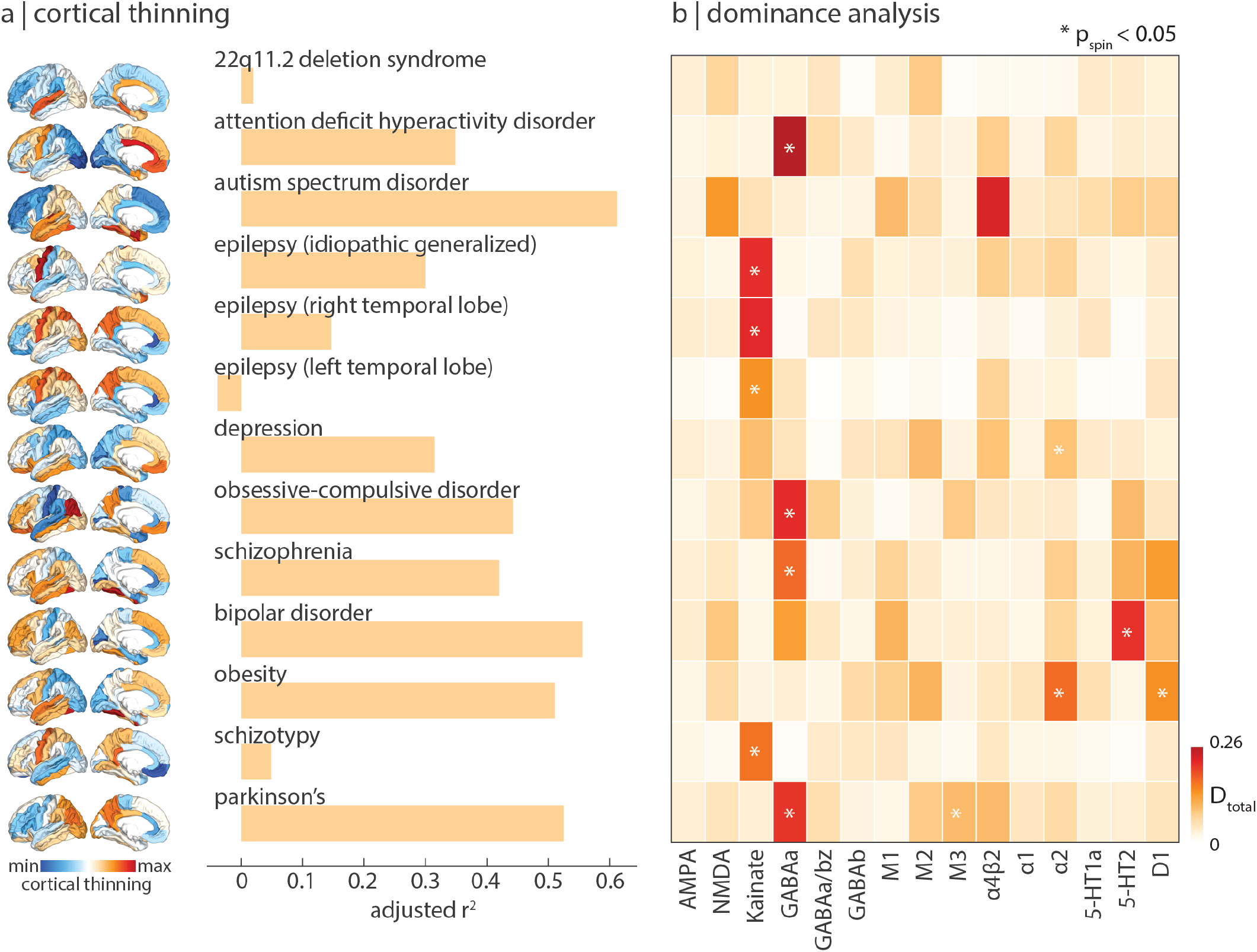
Mapping autoradiography-derived receptors to disease vulnerability. For each disorder, we fit a multilinear regression model between autoradiography-derived receptor densities and cortical thinning, done analogously in Fig. 6. (a) Model fit (adjusted *R*^2^) varies across disorders. (b) Dominance analysis identified the neurotransmitter receptors whose spatial distribution most influence model fit, for each disorder separately. Asterisks indicate significant dominance (*p*_spin_ < 0.05).

**Figure S9.**
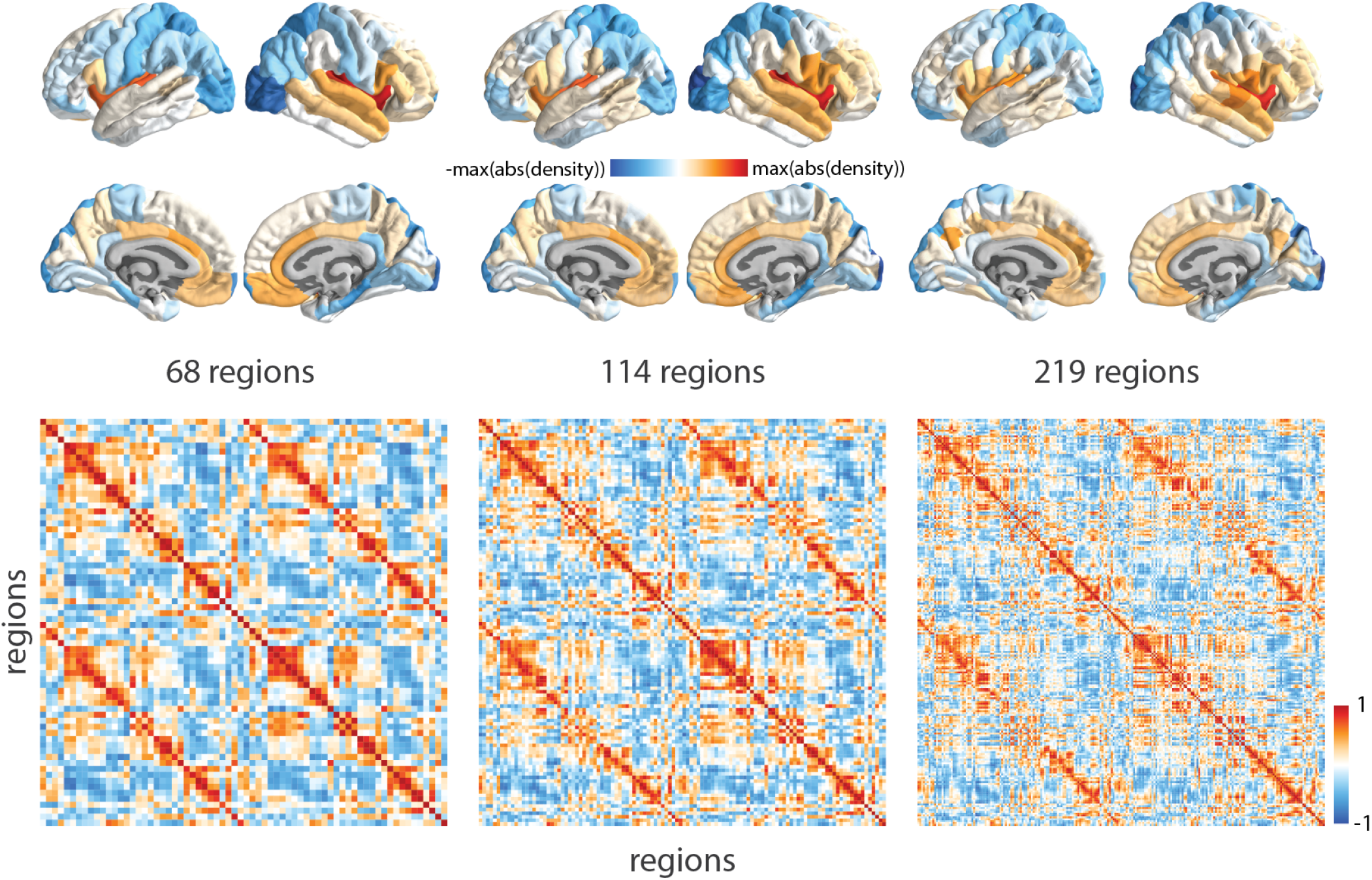
Replicating results using different parcellation resolutions. Top: mean normalized neurotransmitter receptor/transporter density maps are consistent across three increasingly fine parcellation resolutions (68 regions (original), 114 regions, and 219 regions) [18]. Bottom: receptor similarity matrices also demonstrate high conformity across parcellation resolutions.

**Figure S10.**
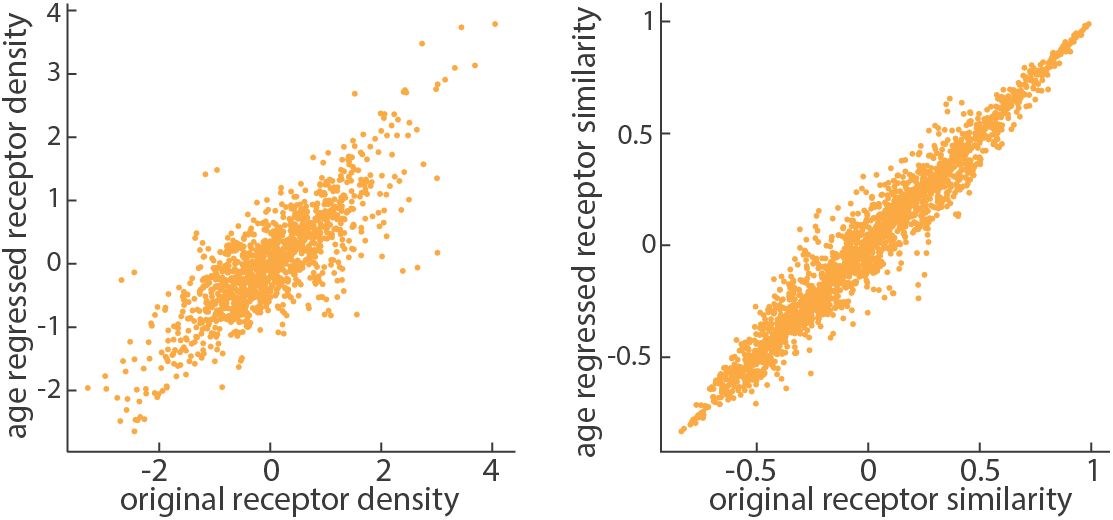
Age has negligible effect on the reported findings. To test age effects of the PET tracer images, we regressed out the relationship between mean age of each tracer map and z-scored receptor densities, at each brain region separately. Age has little impact on receptor density (left; *r* = 0.81) and receptor similarity (right; *r* = 0.98).

**TABLE S2.** Neurosynth terms. Terms that overlapped between the Neurosynth database [151] and the Cognitive Atlas [96] were included in the PLS analysis.

|  |  |  |  |  |
| --- | --- | --- | --- | --- |
| action | eating | insight | naming | semantic memory |
| adaptation | efficiency | integration | navigation | sentence comprehension |
| addiction | effort | intelligence | object recognition | skill |
| anticipation | emotion | intention | pain | sleep |
| anxiety | emotion regulation | interference | perception | social cognition |
| arousal | empathy | judgment | planning | spatial attention |
| association | encoding | knowledge | priming | speech perception |
| attention | episodic memory | language | psychosis | speech production |
| autobiographical memory | expectancy | language comprehension | reading | strategy |
| balance | expertise | learning | reasoning | strength |
| belief | extinction | listening | recall | stress |
| categorization | face recognition | localization | recognition | sustained attention |
| cognitive control | facial expression | loss | rehearsal | task difficulty |
| communication | familiarity | maintenance | reinforcement learning | thought |
| competition | fear | manipulation | response inhibition | uncertainty |
| concept | fixation | meaning | response selection | updating |
| consciousness | focus | memory | retention | utility |
| consolidation | gaze | memory retrieval | retrieval | valence |
| context | goal | mental imagery | reward anticipation | verbal fluency |
| coordination | hyperactivity | monitoring | rhythm | visual attention |
| decision | imagery | mood | risk | visual perception |
| decision making | impulsivity | morphology | rule | word recognition |
| detection | induction | motor control | salience | working memory |
| discrimination | inference | movement | search |  |
| distraction | inhibition | multisensory | selective attention |  |

